# A targeted metabolomic method to detect epigenetically relevant metabolites

**DOI:** 10.1101/2023.11.30.569455

**Authors:** J. Miro-Blanch, A. Junza, J. Capellades, A. Balvay, C. Maudet, M. Kovatcheva, S. Raineri, S. Rabot, J. Mellor, M. Serrano, O. Yanes

## Abstract

Metabolites play a central role in the chemical crosstalk between metabolism and epigenetic marks. Epigenetically relevant metabolites are substrates, products and cofactors that can act as activators or inhibitors of epigenetic enzymes, which control gene expression by adding or removing chemical marks in the DNA, RNA and histones. Diet composition, and biosynthetic pathways encoded in the gut microbiome and the host genome are the main sources of these metabolites for mammals. Despite the increasing interest in the study of the ‘microbiota-nutrient metabolism-host epigenetic axis’ to understand health and disease, there is a lack of a sensitive and easy analytical method to detect epigenetically relevant metabolites simultaneously. Here, we show an straightforward biphasic extraction where the organic phase is directly analyzed by GC-EI MS to detect short-chain fatty acids and formate without chemical derivatization, and the aqueous phase is analyzed by HILIC coupled to ESI-MS/MS, which together can cover >30 epigenetically relevant metabolites in biological samples such as liver, plasma or feces. In addition, we propose a stable isotope tracing method based on multiple-reaction monitoring (MRM) transitions by LC-QqQ MS to understand how ^13^C-labeled glucose or glutamine are used to build SAM and acetyl-CoA, the main methyl and acetyl group donors in epigenetic modifications, respectively. We anticipate that our methods will complement epigenomic and proteomic analyses adding another layer of molecular information towards mechanistic insights.

**Highlights:**

- Host and microbiota metabolites link metabolism with epigenetic regulation.
- Chemical structure diversity in epigenetically relevant metabolites challenges its analysis with a single method.
- A biphasic extraction with no chemical derivatization is able to recover SCFAs and other epigenetically relevant metabolites.
- A novel isotope trace experiment approach allows isotopomer resolution using MS2 data.

## Introduction

Eukaryotes, and especially mammals, have developed a chemical crosstalk between metabolism and the reversible chemical signatures in their DNA, RNA and histone proteins, responding to environmental changes.^1^ Known as epigenetic marks, such signatures can regulate the function, accessibility or 3D architecture of DNA, RNA and histones.^2–7^ The crosstalk between metabolism and epigenetics occurs through specific metabolites that are responsible for regulating the enzymes that add or remove chemical moieties such as methylations, acetylations or glycosylations in these biopolymers.^1^

The host genome encodes enzymes that produce the tricarboxylic acid (TCA) cycle intermediates succinate and alpha-ketoglutaric acid (αKG), that can act as inhibitor and co-substrate of Jumonji demethylases, respectively,^8,9^ or fumarate, capable of modulating histone and DNA methyltransferase activity.^10^ Other examples of host synthesized metabolites are S-adenosylmethionine (SAM), a key metabolite in one carbon metabolism and the main methyl group donor in methylation reactions;^11^ nicotinamide adenine dinucleotide (NAD^+^), which mediates many redox reactions and controls deacetylase activity of sirtuins;^12^ or acetyl coenzyme A (acetyl-CoA), the principal acetyl group donor and the preferred substrate for histone acetyltransferases (HATs).^13^ However, other epigenetically relevant metabolites can only be ingested through the diet or produced by the gut microbiota.^14,15^ This is the case of methionine, an intermediate of the one carbon metabolism involved in the production of SAM, and as an essential amino acid, it needs to be ingested.^14^ In addition to dietary nutrients, the gut microbiota has been recognized as a source of folate, whose levels are associated with global DNA methylation status.^16^ Similarly in its origin, butyric acid is a short-chain fatty acid (SCFA) mainly produced by the gut microbiota that acts as an acetyl group donor and a potent histone deacetylase (HDAC) inhibitor.^15^

Despite the development of specific methods based on LC-MS/MS and GC-MS to detect and quantify epigenetically relevant metabolites individually (e.g, SAM, folate, acetyl-CoA, vitamin B12, butyric acid)^17–21^ or as small sets of related compounds such as folic acid derivatives^22^ or SCFAs^23^, the large structural heterogeneity and physico-chemical diversity of these metabolites has hampered the development of a single and comprehensive method that covers the most important metabolites involved in epigenetic reactions.

Here, we show the optimization of a metabolite extraction protocol with no chemical derivatization combining complementary GC-MS and LC-MS/MS technologies targeting >30 epigenetically relevant metabolites, including intermediates of the TCA cycle^24^, purine and pyrimidine synthesis^25^ vitamins and coenzymes,^13,26^ one carbon metabolism,^27^ essential and non-essential amino acids^28^ and SCFAs.^29^ In addition, we propose a targeted metabolomic approach based on multiple reaction monitoring (MRM) transitions to track the fate of the stable isotope ^13^C from uniformly labeled glucose or glutamine in the synthesis of SAM and acetyl-CoA, the main methyl and acetyl group donors in epigenetic modifications, respectively.

## Results

### Optimizing extraction, chromatographic and mass spectrometry conditions

Due to the chemical heterogeneity of epigenetically relevant metabolites (Supplementary table 1), we explored eight metabolite extraction methods, including biphasic and monophasic extractions (see Methods section for details) coupled to reversed-phase (RP-C18), hydrophilic interaction liquid chromatography (HILIC) or gas chromatography (GC) to improve separation, and MRM transitions for detection by mass spectrometry of these metabolites. Optimization was performed using 42 standard compounds covering the “microbiota-nutrient host metabolism-epigenetic” axis (Figure 1)^30^, which were subsequently monitored in liver tissue and cecal content from mice, and human serum.

**Figure 1.**
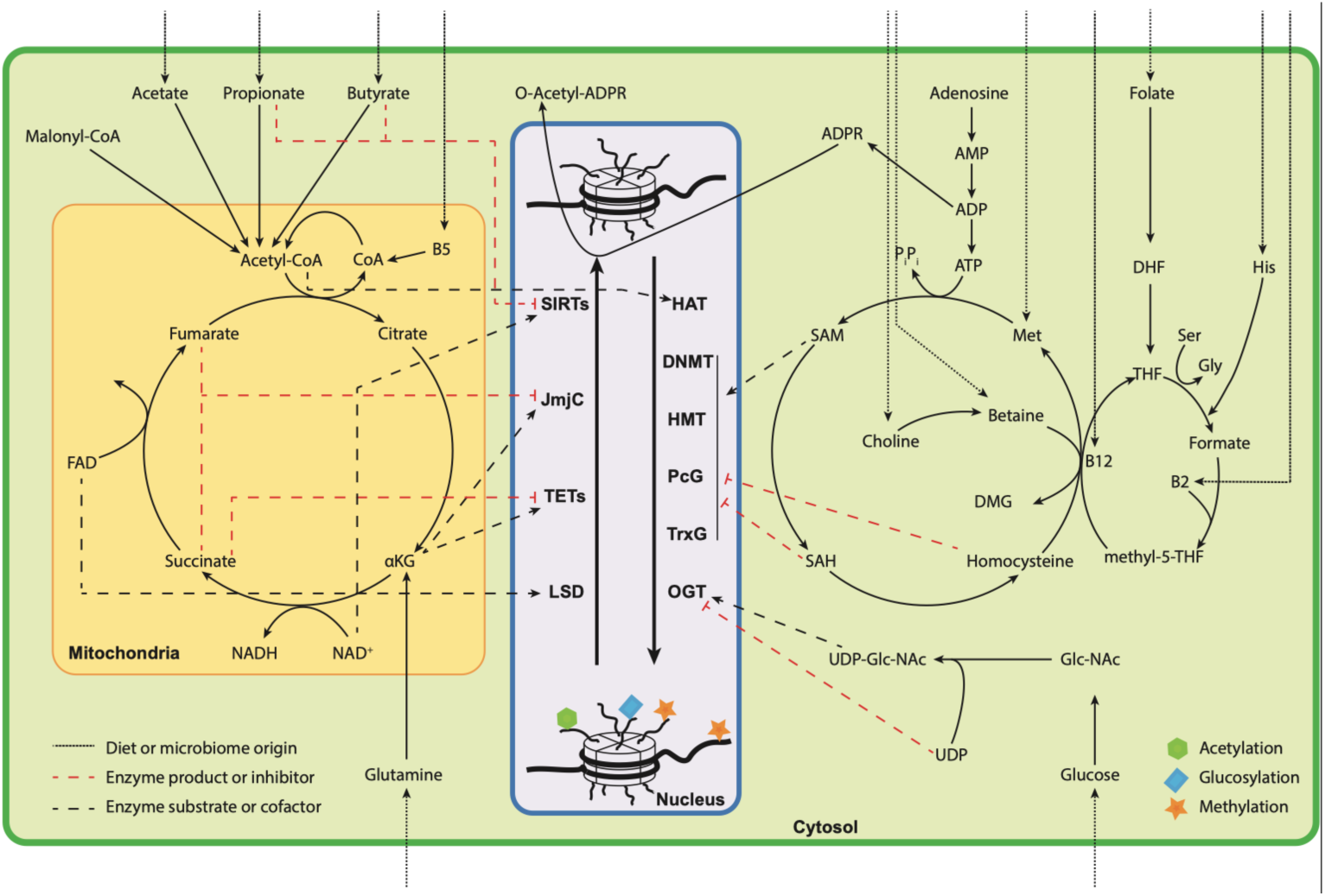
Microbiota-Nutrient host metabolism-epigenetic axis: chemical crosstalk between metabolism and epigenetics. Schematic representation of how epigenetically relevant metabolites regulate epigenetic writers and erasers, connecting microbiota, metabolism and epigenetic marks. CoA: coenzyme A; B5: pantothenic acid; FAD: flavin adenine dinucleotide; ɑKG: alpha-ketoglutarate; NAD+/NADH: nicotinamide adenine dinucleotide/reduced form; AMP: adenosine-5’-monophosphate; ADP: adenosine-5’-diphosphate; ATP: adenosine-5’-triphosphate; ADPR: ADP-ribose; DHF: dihydrofolate; THF: tetrahydrofolate; His: histidine; Ser: serine; Met: methionine; Gly: glycine; DMG: dimethylglycine; SAM: S’-adenosylmethionine; SAH: S’-adenosylhomocysteine; B12: cyanocobalamin; GlcNAc: N’-acetylglucosamine; UDP-GlcNAc: uridine-diphosphate-N-acetylglucosamine; UDP: uridine-diphosphate; SIRTs: sirtuins (deacetylases); JmjC: Jumonji C (demethylase); TETs: ten-eleven translocation methylcytosine dioxygenases (demethylases); LSD: lysine-specific histone demethylases; HAT: histone acetyl transferases; DNMT: DNA methyltransferases; PcG: polycomb group (methylases); TrxG: Trithorax-group proteins (methylases); OGT: O-GlcNAc transferase.

A biphasic extraction adapted from Lotti et al.^23^ consisting in an acidic aqueous solution (with phosphoric acid at 15%) combined with the organic solvent methyl tert-butyl ether (MTBE) resulted in the best extraction method. With the Phospho/MTBE extraction we were able to eficiently extract free SCFAs and formate in the organic phase, while dissolving other relevant epigenetic metabolites in the polar acidic phase. When compared with the best monophasic extraction using ACN/MeOH/H_2_O we observed a high degree of overlap (Figure 2a), however the Phospho/MTBE was the only extraction method capable of recovering SCFAs and formate from every biological matrix tested, that is, liver tissue, caecum and serum (Figure 2b-c,e).

**Figure 2.**
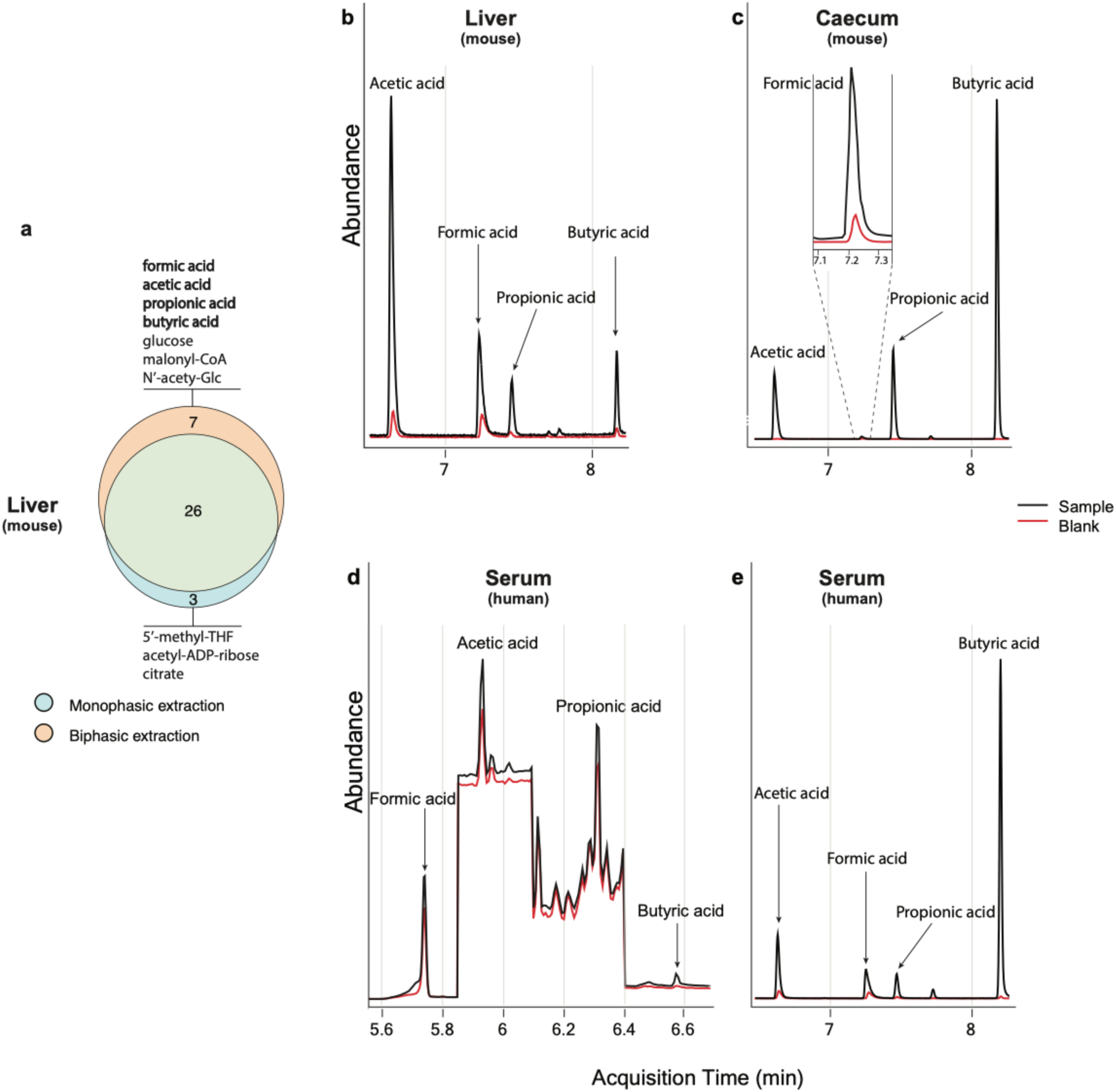
Epigenetically relevant metabolites extracted with a mono- and bi-phasic extraction. (a) Venn diagram showing the overlap of extracted metabolites from liver between the monophasic and biphasic extraction protocols. A complete list of extracted metabolites for each matrix and extraction method can be found in supplementary table 1-2. (b-c) Total ion chromatogram (TIC) of extracted SCFAs and formic acid from mouse liver and caecum content respectively (black line), compared to the corresponding blank sample (red line). (d) TIC of SCFAs and formic acid from human serum using chemical derivatization (black line)^32^ and the corresponding blank sample (red line). (e) TIC of extracted SCFAs and formic acid from human serum using the biphasic extraction without chemical derivatization.

Historically, free SCFAs and formate have been a challenging set of compounds to analyze by liquid and gas chromatography coupled to mass spectrometry, requiring chemical derivatization to stabilize them^15,31^. When we compared the Phospho/MTBE method with another method that required chemical derivatization,^32^ we observed that the derivatization step introduced high background noise in the GC-EI MS spectra, particularly for matrices where the concentration of formate and the SCFAs was low (e.g., serum) (Figure 2d). In contrast, the organic phase of the Phospho/MTBE method analyzed directly by GC-EI MS without chemical derivatization produced cleaner spectra, facilitating the detection and quantification of free SCFAs and formate in liver tissue and serum (Figure 2b,e), where their concentrations are significantly lower than in the gut content (i.e., caecum, figure 2c).^33^

In parallel, we optimized the chromatographic conditions to analyze the acidic aqueous phase of the Phospho/MTBE biphasic extraction. We tested three commonly used stationary phases in metabolomics: (i) reversed phase C18; (ii) hydrophilic interaction liquid chromatographic (HILIC) with ethylene bridged hybrid coated particles (BEH); and (iii) a HILIC zwitterionic stationary phase (HILIC-z). The HILIC-z column performed better than RP-C18 and HILIC, retaining and improving the elution peak shape of most epigenetically relevant metabolites (supplementary figure 1a-d), particularly those phosphorylated, assisted by the additive medronic acid in the mobile phase (supplementary figure 1a-b and d). In addition, we tested another zwitterionic column ZIC-pHILIC (see Methods section), however it resulted in broader peaks by comparing with HILIC-z (supplementary figure 2a-d).

### SCFAs and formic acid measured in germ-free (GF) and conventional (CV) mice

The main advantage of the method presented herein is the ease of measuring SCFAs (acetic, butyric, propionic) and formic acid as well as other epigenetically relevant metabolites with a single metabolite extraction and without using chemical derivatization. To prove that our method can be used to dissect the chemical crosstalk between metabolism and epigenetics –including the gut microbiota metabolic activity– we tested the method in three different matrices: liver and caecum content from germ-free (GF, n=10) and conventional (CV, n=10) mice, and human serum (n=3). GF mice are specially-raised animals devoid of all microorganisms.

As expected, we detected significantly lower levels of SCFAs in the caecum of GF mice compared to the CV group (Figure 3, left panel). In contrast, the amount of SCFA in the liver of GF and CV mice was similar (Figure 3, right panel), which sugest that GF animals compensate for the absence of microbiota by increasing the activity of pathways that generate SCFAs in the liver, such as beta-oxidation, glycolysis or amino acid degradation. Interestingly, free formate levels in caecum exhibit small but significantly lower levels in GF compared to CV mice, indicating that the caecum microbiota is not the major producer of this important metabolite of the one-carbon metabolism.^34,35^

**Figure 3.**
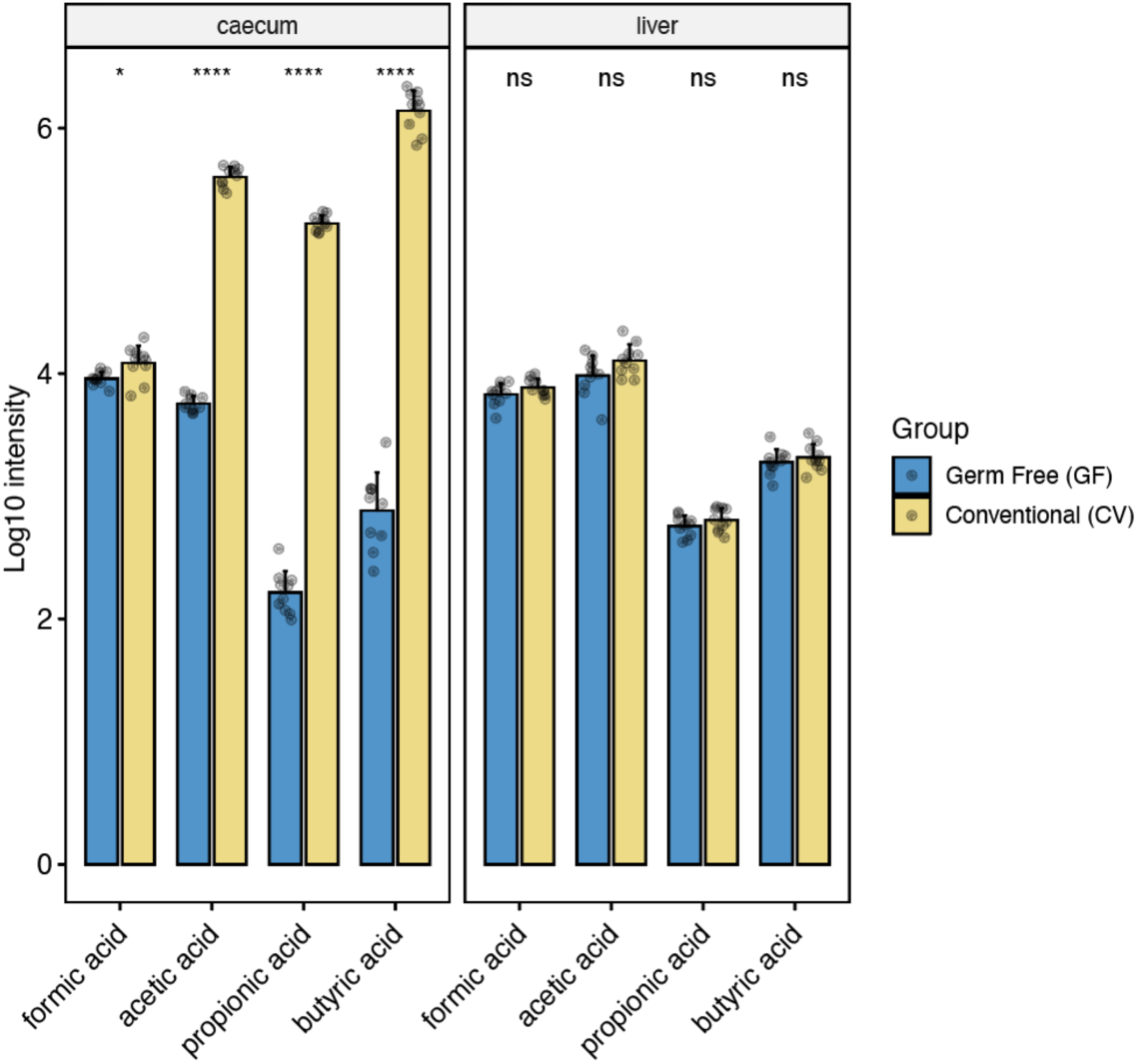
SCFAs and formate in germ-free (GF) and conventional (CV) mice. Abundances in caecum content (left panel) and liver (right panel) of formic acid and short chain fatty acids (SCFAs) in germ-free GF (n=10) and conventional CV (n=10) mice. (Wilcoxon test, after Benjamini-Hochberg (BH) adjusted p-values: ns: p > 0.05; * p <= 0.05; **: p <= 0.01; ***: p <= 0.001; ****: p <= 0.0001)

### Tracking the fate of [U-^13^C]glucose in the synthesis of SAM and acetyl-CoA

The deliberate introduction of a stable isotope into a compound perturbs the isotopologue and isotopomer distribution, and therefore, its mass spectrum. This has been extensively exploited using labeled compounds as tracers for the study of physiological and metabolic processes. However, classical stable isotopic labeling studies are mostly based on MS1 isotopologue distributions,^36^ which only provides information about the number of atoms labeled, lacking positional (i.e., structural) labeling information.

Based on our optimized extraction and chromatographic conditions for epigenetically relevant metabolites, we propose a novel strategy for quantification of positional isotopomers using MRM transitions in QqQ MS (MS2). Our approach feeds from existing information of biosynthetic pathways and the fragmentation pattern of metabolites of interest to design specific MRM transitions that monitor all possible (labeled and unlabeled) isotopomers, determining the exact position and metabolic origin of the labeled ^13^C atoms in the de novo synthesis of SAM and acetyl-CoA (Figure 4 and Supplementary figure 4), the main methyl and acetyl group donors in epigenetic modifications, respectively.

**Figure 4.**
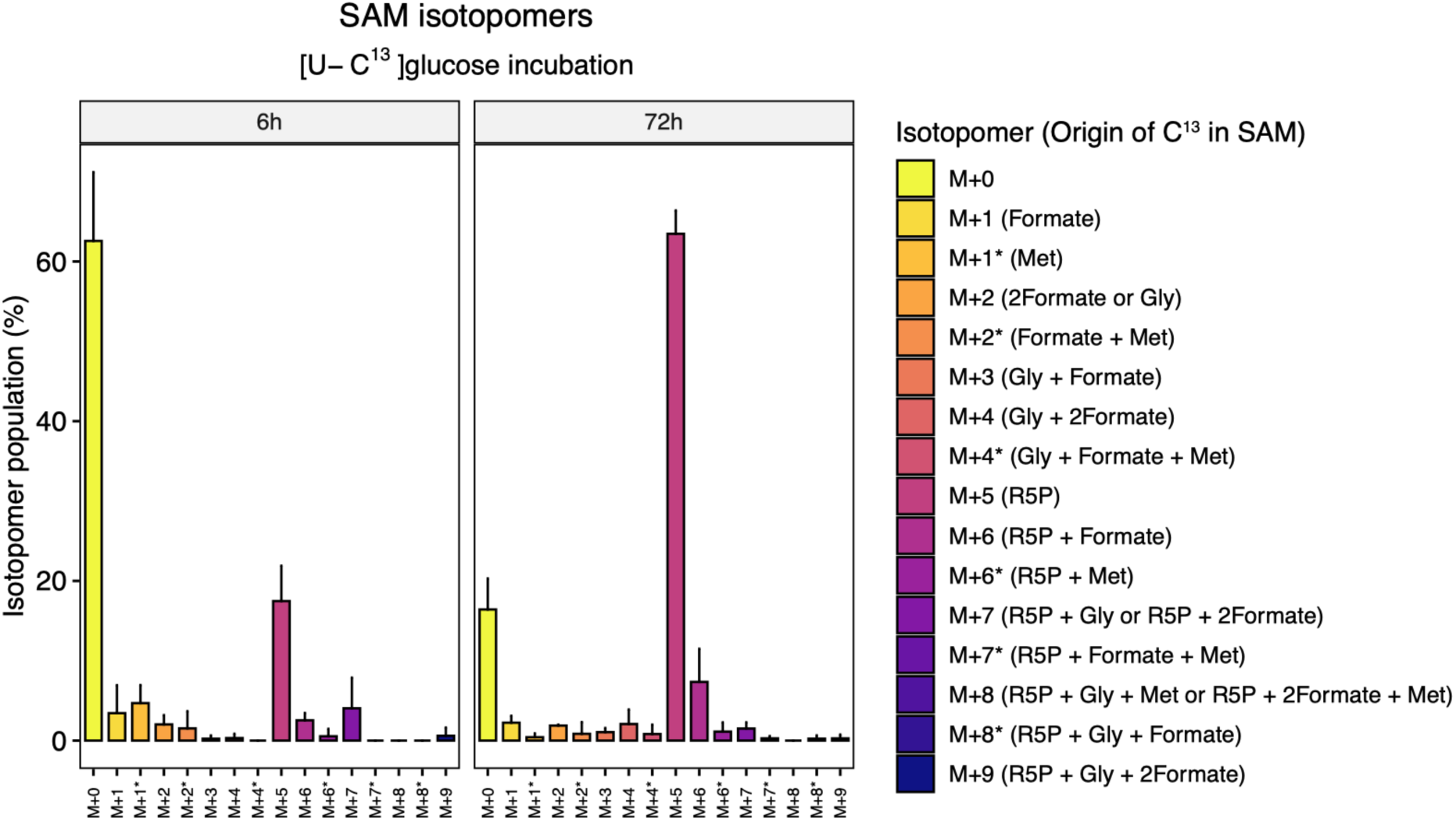
SAM isotopomers distribution after [U-13C]glucose incubation. Barplots representing isotopomers abundance for S-adenosylmethionine (SAM) in an isotope tracing experiment. Embryonic fibroblasts were incubated for 6h and 72h in labeled [U-13C]glucose. The x axis shows the number of extra carbons/neutrons incorporated to the non-labeled compound (M). The colored scale on the right includes the combination of molecules/moieties that give SAM the labeled carbon atoms (legend). Abbreviations: Methionine (Met); Glycine (Gly); Ribose-5-phospate (R5P); * indicates isotopomers with the same number of labeled carbon atoms incorporated, but with different origin.

**Figure 5.**
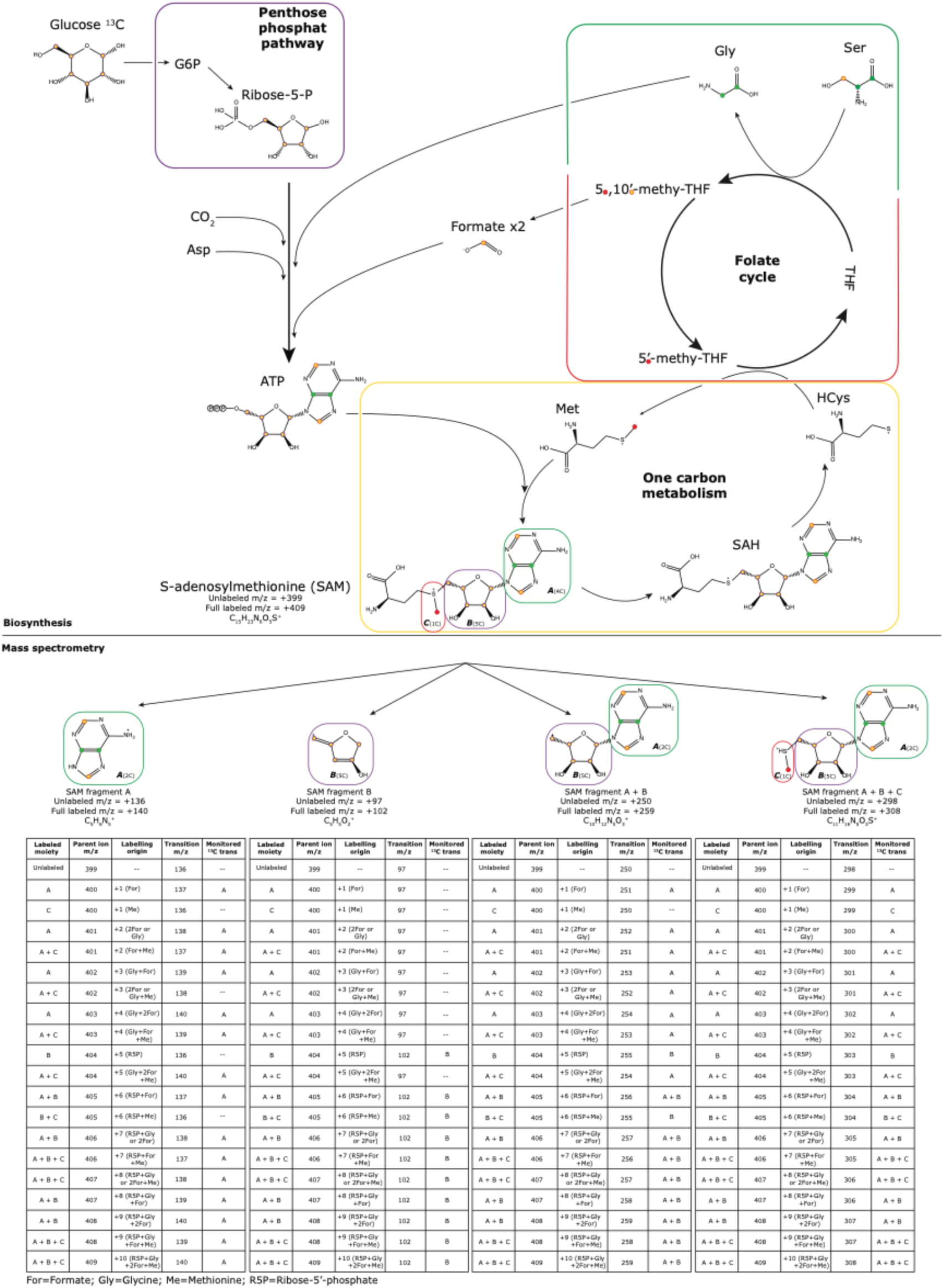
S-Adenosylmethionine (SAM) biosynthesis, labeling and isotopomers. Diagram of metabolic pathways involved in SAM biosynthesis and incorporation of labeled carbon atoms under exposure to [U-^13^C]glucose (biosynthesis). Most intense fragments generated in the mass spectrometer, highlighting the moiety and carbon atoms possibly labeled; and table of parent ions and transitions in the MRM experiment, depicting which moieties get labeled and monitored, and the biosynthetic origin of the labeled atoms (mass spectrometry). The information shown in this table is complementary to the information shown in supplementary table 2.

SAM was analyzed in mouse embryonic fibroblasts (MEF) at two time points (6h and 72h) as described in Kovatcheva M. et al.^37^ Our proposed MRM transitions monitor all biosynthetic combinations contributed by glucose carbons, including the ribose moiety through the pentose phosphate pathway (PPP), the purine nucleotide through de novo synthesis of glycine (C4 and C5) and folate cycle (C2 and C8), and the methyl group from methionine by re-methylation of homocysteine. Our data indicates that glucose metabolism contributes to the synthesis of SAM in a time and moiety dependent manner (Figure 4a-b). After 72h in culture with [U-^13^C]glucose, the most abundant isotopomer corresponds to the ^13^C-enrichment of the ribose moiety (B; M+5). Interestingly, the activity of the folate cycle makes a greater contribution to the de novo purine nucleotide synthesis than the remethylation pathway from homocysteine to methionine and back to SAM. In this latter case, labeling of the methyl moiety (C; M+1) alone, represents 4.6% of the total pool after 6h (after subtracting the natural abundance isotopic distribution). However, the combination of the methyl group (C) with the other two moieties (A and B) reaches 6.6% of the total. This sugests that a non-negligible fraction of glucose-derived one-carbon units (through the glycine and serine pathway) serve as methyl donors for the re-methylation of homocysteine to support the methionine cycle,^37^ in contrast to recent observations in activated macrophages.^38^ Overall, our approach shows the high number of different labeled isotopomers in SAM and their percentages within the total pool. The proposed method determines the exact position and biosynthetic origin of the labeled atoms, which would not be distinguished by analyzing isotopologue information from MS1 data.

Acetyl-CoA was analyzed in U-87 MG cells isolated from malignant gliomas. Acetyl-CoA has three chemical moieties originating from glucose metabolism (Supplementary figure 3): the purine nucleotide through de novo synthesis of glycine (C4 and C5) and folate cycle (C2 and C8); the ribose from the pentose phosphate pathway; and the acetyl group from pyruvate through glycolysis (named A, B and C respectively in supplementary figure 3). Similar to SAM, our data indicates that carbon atoms from glucose metabolism contribute to the synthesis of acetyl-CoA in a time and moiety dependent manner (Supplementary figure 4). As in the previous case, after 72h of [U-^13^C]glucose in the culture media, the ribose moiety (B; M+5) is the predominant isotopomer, representing >50% of the total pool, which sugests an active metabolic Aux through the pentose-phosphate pathway (PPP).^39^ The second most abundant isotopomer from [U-^13^C]glucose is the acetyl moiety (C; M+2), particularly at 20h (20-25% of the total pool) (Supplementary figure 4). Glutamine carbons, in contrast, are mainly used to build the acetyl moiety (C; M+2), either from the oxidative or the reductive carboxylation of glutamine when entering the TCA cycle through alpha-ketoglutarate. Yet, the contribution of [U-^13^C]glutamine to the acetyl group only represents ∼10% of the analyzed isotopomers at the two time points studied, indicating a low but steady flow of glutamine carbons to acetyl-CoA. Similar to SAM, the combination of different moieties (e.g., AB, BC, ABC) results in a less abundant but non-negligible fraction of isotopomers.

## Material and methods

### Standards and metabolomic reagents

Methanol (MS grade), acetonitrile (ACN, MS grade), pyridine, methoxyamine hydrochloride (MA), formic acid (MS grade), N-methyl-N-(trimethylsilyl)-trifluoroacetamide (TMS) and ammonium acetate were purchased from Thermo Fisher Scientific (Waltham). All pure standards in supplementary table 1, 2,3,4,5,6-Pentafluorobenzyl bromide (PFBBr), hexane and acetone were purchased from Sigma-Aldrich (St. Louis). Acetyl-ADP-ribose was purchased from Santa Cruz Biotechnology (Dallas). Medronic acid was purchased from Agilent Technologies (Santa Clara).

### Germ-free and conventional mice samples

Liver and cecal content were obtained from nine-week-old female germ-free (GF) and conventional (CV) C57BL/J6 mice strain. The GF mice were procured by the breeding unit Anaxem (INRAE, Jouy-en-Josas, France; Anaxem license number: B78-33-6). The CV mice were purchased from Charles River Laboratories (L’Arbresle, France) and kept in the Anaxem facilities. The GF mice were housed in sterile isolators (Getinge, Les Ulis, France) in individual cages. Fresh defecations were used to ensure sterile conditions weekly by microscopic examination and screening cultures. CV mice were housed in the same type of isolators but non-sterile, to ensure the same environment and stress between experimental groups. In each isolator, mice were kept in enriched home cages containing paper towels, wooden sticks and sterile bedding made of wood shavings, and free access to autoclaved tap water and γ-irradiated (45 kGy) standard chow diet (R03; Scientific Animal Food and Engineering, Augy, France). The experimental procedures were performed in accordance with European guidelines for the care and use of laboratory animals and approved by the ethics committee of the INRAE Research Centre at Jouy-en-Josas (approval reference: 17-14). Ten mice females were used for each microbiota status, with a total n=20 mice. All mice were housed individually for measurement of individual food consumption. Body weight and food were weighted once a week from the age of 4 to 9 weeks. And animal room temperature was maintained at 20–24°C with a strict 12-h light/dark cycle (lights open at 7:30 am).

### Cell culture samples in ^13^C media

#### For the analysis of SAM

Doxycycline-inducible i4F MEFs were cultured as described in Kovatcheva M. et al. ^37^, with 1 mg ml_−1_ doxycycline. At 72 h after the addition of doxycycline, cells were transferred to complete KSR media containing a final concentration of 4500 mg/L [U-_13_C]glucose 99% (CAT# CLM-1396-1, Sigma-Aldrich). This is the same concentration of unlabelled glucose normally found in the complete KSR media, and was generated by ordering custom, glucose-free DMEM (Life Technologies, ME22802L1). Six hours after the addition of labeled media, a subset of wells was harvested by scraping in PBS and centrifugation (300g for 5 min); supernatant was removed and pellets were snap frozen. At 72 h after the addition of the labeled media (that is, 6 days into reprogramming), a subset of wells was harvested by scraping in PBS and centrifugation (300g for 5 min); supernatant was removed and pellets were snap frozen.

#### For the analysis of acetyl-CoA

Brain immortalized cell line U-87 MG (ATCC® HTB-14™) (WT) cells were plated in a density of 1 million per T75 plate in 10 mL complete ATCC EMEM 30-2003 medium and have 5 replicates per condition. Two conditions were tested,^13^C labeled [U-^13^C]glucose and labeled [U-^13^C]glutamine. After 8 h, the medium was changed to [U-^13^C]glucose, add 10 mL per T75 (Nacalai Tesque 09848 + [U-^13^C]glucose + Sodium Pyruvate + FBS); or ^1^[U-^13^C]glutamine, add 10 mL per T75 (Sigma M5650 + [U-^13^C]glutamine + Sodium Pyruvate + FBS). A complete description of medium composition can be found in the supplementary table 3.

### Metabolite extraction methods on mice samples

Frozen lyophilized and pulverized mice liver was used for optimizing and choosing the best metabolite extraction method. Two different types of extractions were tested: 1) monophasic extractions, using one or more miscible solvents and having a homogeneous mixture; and 2) biphasic extractions, using two or more solvents forming two immiscible phases. Once the final extraction method was chosen, we tested the protocol in two additional matrices: mouse caecum content and human serum.

### Monophasic extractions

#### ACN/MeOH/H_2_O

A volume of 400 μL of acetonitrile methanol water solution (ACN:MeOH:H_2_O in a 4:4:2 volume proportion) was added to 5 mg of pulverized liver tissue and vortex for 1 min. Here, 3 different pH (pH = *3.2*, *6.6* and *9.1*) of the extraction solution were tested. The sample was incubated in liquid nitrogen for 30 s, sonicated for 20-30 s in a water bath and vortexed for 30 s, repeating these steps up to a total of three times. Next, the sample was incubated at −20°C for 60 min and then centrifuged 10 min at 22000 g (4°C). Finallys 100 μL of supernatant was transferred into HPLC vial insert for LC-MS analysis, results are shown in supplementary figure 5.

#### MeOH/H_2_O (8:2 and 1:1)

A volume of 500 μL of methanol water (MeOH/ H_2_O, in a 8:2 or 1:1 volume proportion) solution was added to 5 mg of pulverized liver tissue, vortexed for 1 min followed by 3 cycles of incubation in liquid nitrogen for 30 s, sonicated for 30 s in a water bath and vortexed for 30 s. The extraction was incubated on ice for 60 min to allow proteins to precipitate and then centrifuged 10 min at 22000 g (4°C), 50 μL of supernatant was transferred into a HPLC vial insert for LC-MS injection.

#### Metaphosphoric acid (MPA)

A volume of 300 μL of cold acetonitrile water (ACN/H_2_O, 1/1) solution with 1% of metaphosphoric acid was added to 5 mg of pulverized liver tissue, vortexed for 30 s, followed by 3 cycles of incubation in liquid nitrogen for 30 s, sonicated for 30 s in a water bath and vortexed for 30 s. After 2 hours incubation at −20°C the samples were centrifuged 10 min at 22000 g (4°C), and 100 μL of supernatant was transferred to a HPLC vial insert.

#### •2-Mercaptoethanol (BME)

A volume of 500 μL of 50 mM phosphate buffer (1x), at pH=7, with 1 % of ascorbic acid and 0.1 % 2-mercaptoethanol was added to 5 mg of pulverized liver tissue, vortexed for 1 min, followed by 3 cycles of incubation in liquid nitrogen for 30 s, sonicated for 30 s in a water bath and vortexed for 30 s. After 60 min incubation on ice, the sample was centrifuged 10 min at 22000 g (4°C), and 100 μL of supernatant transferred to an HPLC vial insert.

### Biphasic extractions

#### H2O phosphoric 15% solution: Methyl Tert butyl Ether (Phospho/MTBE)

A volume of 200 μL of water with 15% of phosphoric acid was added to 5 mg of pulverized liver tissue and vortex for 1 min, followed by 3 cycles of incubation in liquid nitrogen for 30 s, sonicated for 30 s in a water bath and vortexed for 30 s. Then 200 μL of methyl tert butyl ether (MTBE) was added and vortexed for 1 min. After 60 min incubation on ice the sample was centrifuged 10 min at 22000g (4 °C) and formed two separated phases. The acidic H_2_O was the bottom phase and MTBE was the top phase. 100 μL of the acidic H_2_O phase were diluted in 900 μL of pure acetonitrile (ACN) followed by 1h incubation at −20°C to facilitate protein precipitation. Then the samples were centrifuged for 10 min at 22000 g (4 °C) and 100 μL of supernatant (bottom phase) was transferred to an HPLC vial, ready for LC-MS analysis. 100 μL of MTBE (top phase) was directly transferred after the first centrifugation to an HPLC vial insert for direct GC-MS analysis.

#### H_2_O formic 0.1 %/Diethyl Etether (H_2_O/DEE)

A volume of 400 μL of water with 0.1 % of formic acid was added to 5 mg of pulverized liver tissue and vortex for 1 min, followed by 3 cycles of incubation in liquid nitrogen for 30 s, sonicated for 30 s in a water bath and vortexed for 30 s. Then 400 μL of diethyl ether (DEE) was added and vortexed for 1 min. After 60 min incubation on ice the sample was centrifuged 10 min at 22000 g (4 °C) and formed two separated phases. Water (H_2_O) was the bottom phase and diethyl ether (DEE) was the top phase, from which 400 μL of each phase were separated in different HPLC vials. For LC-MS analysis, an aliquot of 50 μL of aqueous (bottom) phase was transferred to a HPLC vial insert for direct LC-MS analysis. For GC-MS analysis, both phases (H_2_O and DEE) were analyzed with the following downstream preparation: H_2_O phase was frozen at −80 °C followed by 2 or more hours in the lyophilizator until dry; and DEE phase was dried under N_2_ flux. Lyophilized or dry samples were derivatized as described here. Chemical derivatization. Metabolite standards for method optimization or liver extracts to be analyzed by the GC-MS method were chemically derivatized as follow: 1) adding 40 μL of methoxyamine in pyridine (30 μg/μL) and incubated for 45 minutes at 60°C; and 2) adding 25 μL of N-methyl-N-trimethylsilyltrifluoroacetamide (TMS) with 1% trimethylchlorosilane (Thermo Fisher Scientific) and incubated for 30 minutes at 60°C. The 65 μL of derivatized sample extract or metabolite standards were transferred into HPLC vial inserts for GC-MS analysis.

#### H_2_O formic 0.1 %/Methanol/Diethyl Ether (Mix)

This extraction was a mix between the H_2_O/DEE extraction and the MeOH/H_2_O (1/1) with 0.1 % of formic acid extraction. A volume of 500 μL of MeOH/H_2_O (1/1) extraction with 0.1 % of formic acid were added to the sample, followed by the 3 cycles of incubation in liquid nitrogen for 30 s, sonicated for 30 s in a water bath and vortexed for 30 s. Then 500 μL of DEE was added and vortexed for 1 min. After 60 min incubation on ice, the sample was centrifuged 10 min at 22000 g (4 °C) and formed two separated phases. Water (H_2_O) was the bottom phase and diethyl ether (DEE) was the top phase, 50 μL of the water phase were separated for LC-MS analysis. The upper phase (DEE) was separated in a different HPLC vial to be dried under N_2_ Aow and derivatizated as described in the H_2_O/DEE extraction.

#### 2,3,4,5,6-Pentafluorobenzyl bromide (PFBBr)

A volume of 20 μL of phosphate buffer (0.5M pH 8.0) was added to 50 μL of human serum. Add 130 μL of PFBBr solution (100 mM in acetone) and vortex for 1 min followed by an incubation of 15 min at 60 °C. After the incubation, wait 2-3 min at RT to cold down the solution and add 330 μL of hexane and vigorously vortex. After vortexing two phases are clearly separated, corresponding the upper phase to the organic solvent containing SCFAs. 100 μL of the supernatants was transferred to a GC vial for analysis.

### LC-MS/MS analysis of epigenetically relevant metabolites

#### On standards and test samples

LC-MS method optimization was performed using around 1ppm dilutions of pure metabolites standards in table 1, except for the SCFAs. Pure metabolite standards and sample extracts were analyzed by LC-MS. Metabolites in the sample extractor mixture of pure standards were separated using a 1290 Infinity Series liquid chromatograph (Agilent Technologies), coupled with positive and negative electrospray ionization (-ESI & +ESI) at the same time ^40^, followed by spectral mass data acquisition with a 6490 triple quadrupole QqQ mass spectrometer (Agilent Technologies). The mass spectrometer was operated in multiple reaction monitoring (MRM), and samples for LC-MS analysis were not derivatized. Extracted metabolites or mixtures of pure standards were kept in vials at −80°C until the LC-MS analysis was performed. Standard mixtures and metabolite extractions were separated using one of the following chromatographic columns: Luna Omega 1.6 μm Polar C18 100 Å, LC Column 100 x 2.1 mm (Phenomenex) as reverse phase column; and ACQ_UITY UPLC BEH HILIC Column, 130Å, 1.7 µm, 2.1 mm X 150 mm (Waters), InfinityLab Poroshell 120 HILIC-Z (HILIC-Z), 2.1 x 100 mm, 2.7 μm (PEEK lined) (Agilent Technologies) or the SeQ_uant® ZIC-pHILIC (pHILIC) 5µm polymer 150 x mm (Merck) as hydrophilic interactions chromatography (HILIC) columns. The HILIC-Z and SeQ_uant columns have a zwitterionic stationary phase, having at the same time positive and negative charges, allowing the retention of challenging polar and charged metabolites and with a biger dynamic range. All columns were operated with their correspondent precolumn.

#### On experimental samples

Metabolites in liver extracts were separated by using either the HILIC-Z or the pHILIC columns, using either 50 mM ammonium acetate with 5 μM of medronic acid solution or a 20 mM ammonium acetate with 2.5 μM of medronic acid solution, respectively as mobile phase A and 100% acetonitrile (ACN) as phase B. For the HILIC-Z, the mobile phase flux was set to 0.4 mL/min with a linear gradient elution that started at 98% B (time 0-2 min), followed by an isocratic gradient from 98% and finishing at 40 % B (time 2-9 min), then back to 98% B (time 9-9.5 min) and a holding time 3 min and a half at 98% B (time 9.5-13 min) to allow system stabilization. For the pHILIC, the mobile phase flux was set to 0.2 mL/min with a linear gradient elution that started at 85% B (time 0-2 min), followed by an isocratic gradient from 85% and finishing at 40 % B (time 2-12 min), then back to 85% B (time 12-12.5 min) and a holding time 6 min and a half at 85% B (time 12.5-19 min) to allow system stabilization. Ions were generated using positive and negative electrospray ionization (+ESI & -ESI) and spectral data measured with a 6490 QqQ mass spectrometer (Agilent Technologies) operated in both positive and negative ion mode. The injection volume was set to 3 μL. The mass spectrometer parameters were: drying and sheath gas temperatures 270°C and 400°C, respectively; source and sheath gas flows 15 and 11 L/min, respectively; nebulizer flow was set to 35 psi; positive and negative capillary voltage were both set at 3000V; nozzle voltages were 1000V and −1500V, in positive and negative respectively; and iFunnel in positive HRF and LRF 130 and 100V, respectively; and iFunnel in negative HRF and LRF 110 and 60V, respectively. The ions and transitions that have been monitored can be found in supplementary *table 1* in the parent m/z, 1^st^ and 2^nd^ m/z transitions columns, together with each collision energy (CE) used. We manually quantified all pure standards (for method optimization) and metabolite extraction peaks from samples with the Q_ualitative Analysis of MassHunter Workstation (Agilent Technologies).

### GC-MS/MS analysis of SCFAs

#### On standards and test samples

GC-MS method optimization was performed by using around 1ppm dilutions of pure standards of derivatized and non-derivatized formic, acetic, propionic and butyric acid (SCFAs, in table 1). Pure metabolite standards and sample extracts analyses performed by GC-MS, were separated using a 7890A gas chromatograph (Agilent Technologies), coupled with one of the two different ionization sources tested: electron impact (EI) or chemical ionization (CI), followed by spectral mass data acquisition with a 7000 QqQ mass spectrometer (Agilent Technologies). The mass spectrometer was operated in multiple reaction monitoring (MRM) for derivatized and non-derivatized samples. Derivatized extracts or standards were injected (1μL) into the gas chromatograph with a split inlet and a J&W Scientific DB5−MS+DG (5MS) stationary phase column of 30 m × 0.25 mm i.d., 0.1 μm film (Agilent Technologies). Non-derivatized samples or pure standards analyzed were injected (1μL) into the gas chromatograph system with a split inlet in a J&W Scientific HP-FFAP (FFAP) stationary phase column 30m × 0.25mm i.d., 0.25 μm film (Agilent Technologies).

#### On samples

The extracted liver SCFAs in the organic phase (top) of the Phospho/MTBE extraction were separated by using a J&W Scientific HP-FFAP (FFAP) stationary phase column (30m × 0.25mm i.d., 0.25 μm film, Agilent Technologies) when samples were non-derivatized and into a J&W Scientific DB5−MS+DG (5MS) stationary phase column (30 m × 0.25 mm i.d., 0.1 μm film, Agilent Technologies) when samples were derivatized. Sample metabolites were separated with a 7890A gas chromatograph (Agilent Technologies), coupled with an electron impact (EI) ionization source, followed by spectral mass data acquisition with a 7000 QqQ mass spectrometer (Agilent Technologies). The GC-MS/MS conditions for the derivatized samples were as follow: the 5MS column was used, carrier gas was helium at 7.5 mL/min flow, split ratio was set to 2:1, oven temperature was set to 50 °C for 2 min, followed by a ramp of 30 °C/min up to 220 °C and holding this temperature for 3 min, source heather was set to 300 °C and ionization was achieved by electron impact (EI) at 70 eV. The GC-MS conditions for the non-derivatized samples were as follow: the FFAP column was used, carrier gas was helium at 54.5.1 mL/min flow, split ratio was set to 10:1, oven temperature was set initially to 40 °C for 0 min, followed by a ramp of 12 °C/min up to 130 °C, then a ramp of 30 °C/min up to 250 °C and holding this temperature for 5 min, source heather was set to 250 °C and ionization was achieved by electron impact (EI) at 70 eV. We manually quantified all metabolite extraction peaks with the Q_ualitative Analysis of MassHunter Workstation (Agilent Technologies).

#### 13C flux analysis from labeled glucose and glutamine to acetyl-CoA

Metabolomics analyses were performed on cultured brain cell line U-87 MG with ^13^C media as follows: at 20h and 72h media was aspirated and washed one time with cold PBS (phosphate buffer or any other physiological buffer) to avoid carry over from media components. In order to extract the metabolites from the cells, 1mL of extraction solution (80:20 methanol:water) prechilled at −80°C was added to the cells and incubated for 15 min at −80°C to break the cells. After, the cells were scraped off the dish with a cell scraper and the suspension (cells in the extraction solution) transferred into a clean 1.5 mL, followed by 10 min centrifugation at 2000g at 4°C to pellet cellular debris. Supernatant was transferred into a new 1.5 mL tube and set aside on ice. The metabolite extraction was repeated two more rounds on the cell pellets, using 125 μL of extraction solution. The corresponding supernatants were combined with the supernatant of the first round of extraction in the plate and stored at −80°C. Supernatants were dried under N_2_ gas flow until completely dry, in order to resuspend them in the same volume. The dried extracts were resuspended in 400 μL of extraction solution (80:20 methanol:water) and 100 μL were put in a HPLC vial for targeted analysis.

The extracted metabolites were separated using an InfinityLab Poroshell 120 HILIC-Z (Agilent Technologies) column, using a 50 mM ammonium acetate with 5 μM of medronic acid as phase A and 100% acetonitrile (ACN) as phase B. The mobile phase flux was set to 0.4 mL/min with a linear gradient elution that started at 98% B (time 0-2 min), followed by an isocratic gradient from 98% and finishing at 40 % B (time 2-9 min), then back to 98% B (time 9-9.5 min) and a holding time 3 min and a half at 98% B (time 9.5-13 min) to allow system stabilization. Ions were generated using electrospray ionization (ESI) and spectral data measured with a 6490 QqQ mass spectrometer (Agilent Technologies) operated in both positive and negative modes. The injection volume was 5 μL. The mass spectrometer parameters were the same as described in the LC-MS/SM methods section. The parent ions, transitions and collision energies that have been monitored can be found in supplementary table 2. We calculated most relevant and fully labeled isotopomers for acetyl-CoA based on: the total number of carbon atoms possibly labeled; pathways involved in the synthesis of their moieties from labeled glucose or glutamine; and previous knowledge of fragmentation patterns generated by MS analysis of the unlabelled metabolites. All samples were manually quantified by extracting and integrating the MRM peak intensities with Q_ualitative Analysis of MassHunter Workstation (Agilent Technologies).

#### 13C flux analysis from glucose to SAM

Samples were analyzed with an UHPLC 1290 Infinity II Series coupled to a QqQ/MS 6490 Series from Agilent Technologies (Agilent Technologies). The source parameters applied operating in positive electrospray ionization (ESI) were gas temperature: 270 °C; gas flow: 15 l min^−1^; nebulizer: 35 psi; sheath gas heater, 400 a.u.; sheath gas flow, 11 a.u.; capillary, 3000 V; nozzle voltage: 1000 V.

The chromatographic separation was performed with an InfinityLab Poroshell 120 HILIC-Z 2.7 μm, 2.1 mm × 100 mm column (Agilent Technologies), starting with 90% B for 2 min, 50% B from minute 2 to 6, and 90% B from minute 7 to 7.2. Mobile phase A was 50 mM ammonium acetate in water, and mobile phase B was acetonitrile. The column temperature was set at 25 °C and the injection volume was 2 μl. MRM transitions and collision energies for SAM isotopomers are detailed in supplementary table 2.

