## Supplementary Materials for "A targeted metabolomic method to detect epigenetically relevant metabolites"

Title:


**Highlights:**

- Host and microbiota metabolites link metabolism with epigenetic regulation.
- Chemical structure diversity in epigenetically relevant metabolites challenges its analysis with a single method.
- A biphasic extraction with no chemical derivatization is able to recover SCFAs and other epigenetically relevant metabolites.
- A novel isotope trace experiment approach allows isotopomer resolution using MS2 data.

### Abstract

Metabolites play a central role in the chemical crosstalk between metabolism and epigenetic marks. Epigenetically relevant metabolites are substrates, products and cofactors that can act as activators or inhibitors of epigenetic enzymes, which control gene expression by adding or removing chemical marks in the DNA, RNA and histones. Diet composition, and biosynthetic pathways encoded in the gut microbiome and the host genome are the main sources of these metabolites for mammals. Despite the increasing interest in the study of the 'microbiota-nutrient metabolism-host epigenetic axis' to understand health and disease, there is a lack of a sensitive and easy analytical method to detect epigenetically relevant metabolites simultaneously. Here, we show an straightforward biphasic extraction where the organic phase is directly analyzed by GC-EI MS to detect short-chain fatty acids and formate without chemical derivatization, and the aqueous phase is analyzed by HILIC coupled to ESI-MS/MS, which together can cover >30 epigenetically relevant metabolites in biological samples such as liver, plasma or feces. In addition, we propose a stable isotope tracing method based on multiple-reaction monitoring (MRM) transitions by LC-QqQ MS to understand how $^{13}C$-labeled glucose or glutamine are used to build SAM and acetyl-CoA, the main methyl and acetyl group donors in epigenetic modifications, respectively. We anticipate that our methods will complement epigenomic and proteomic analyses adding another layer of molecular information towards mechanistic insights.

### Supplementary material

#### Supplementary figures:

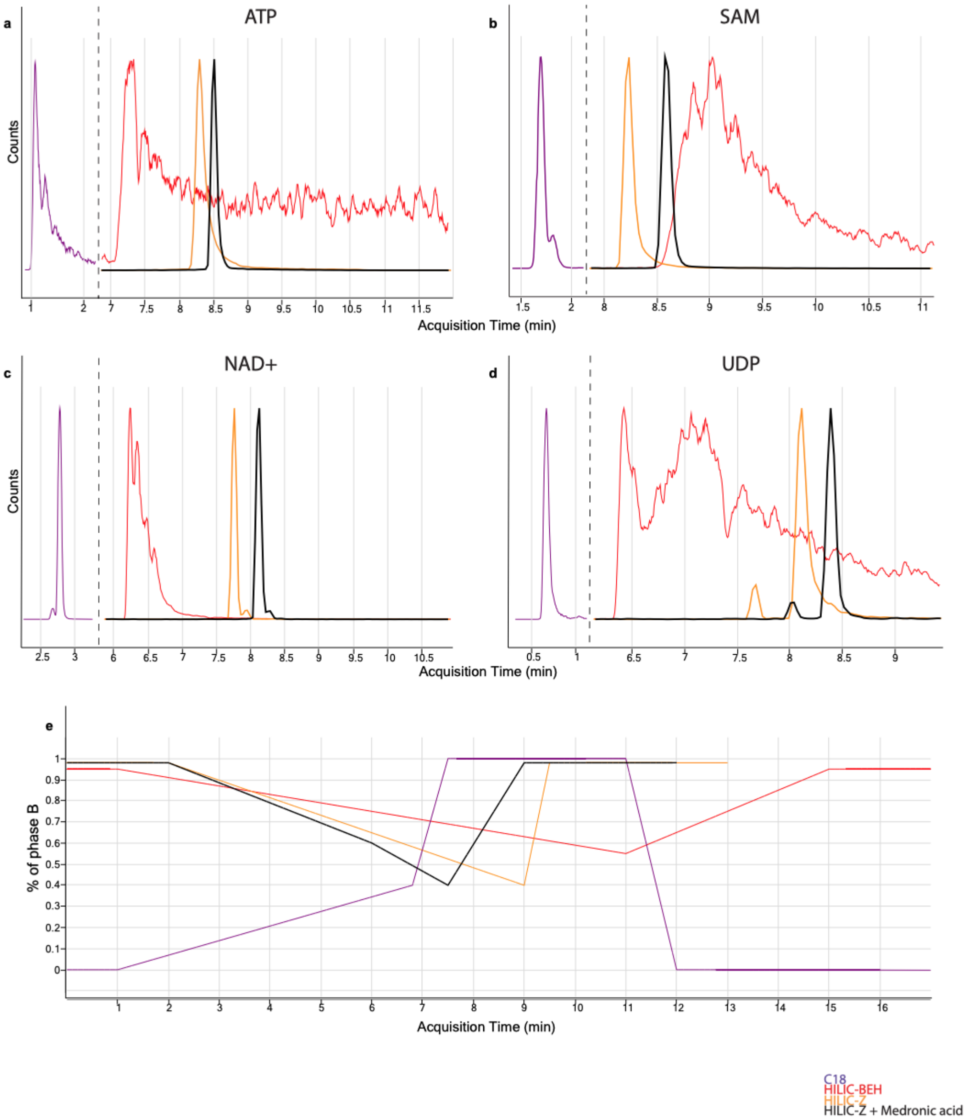


**Supplementary figure 1.** Positively charged metabolites or with a phosphate group dictate LC conditions.

(a-d) Multiple reaction monitoring (MRM) extracted chromatograms of ATP, SAM, NAD+ and UDP compared across four different chromatographic conditions. (e) Chromatographic elution profile showing the percentage of B (organic) phase. C18: reverse phase C18 column; HILIC-BEH: hydrophilic interactions liquid chromatographic (HILIC) ethylene bridged hybrid (BEH); HILIC-Z: HILIC zwitterionic column; HILIC-Z + medronic: HILIC-Z column run with medronic acid in the aqueous phase.

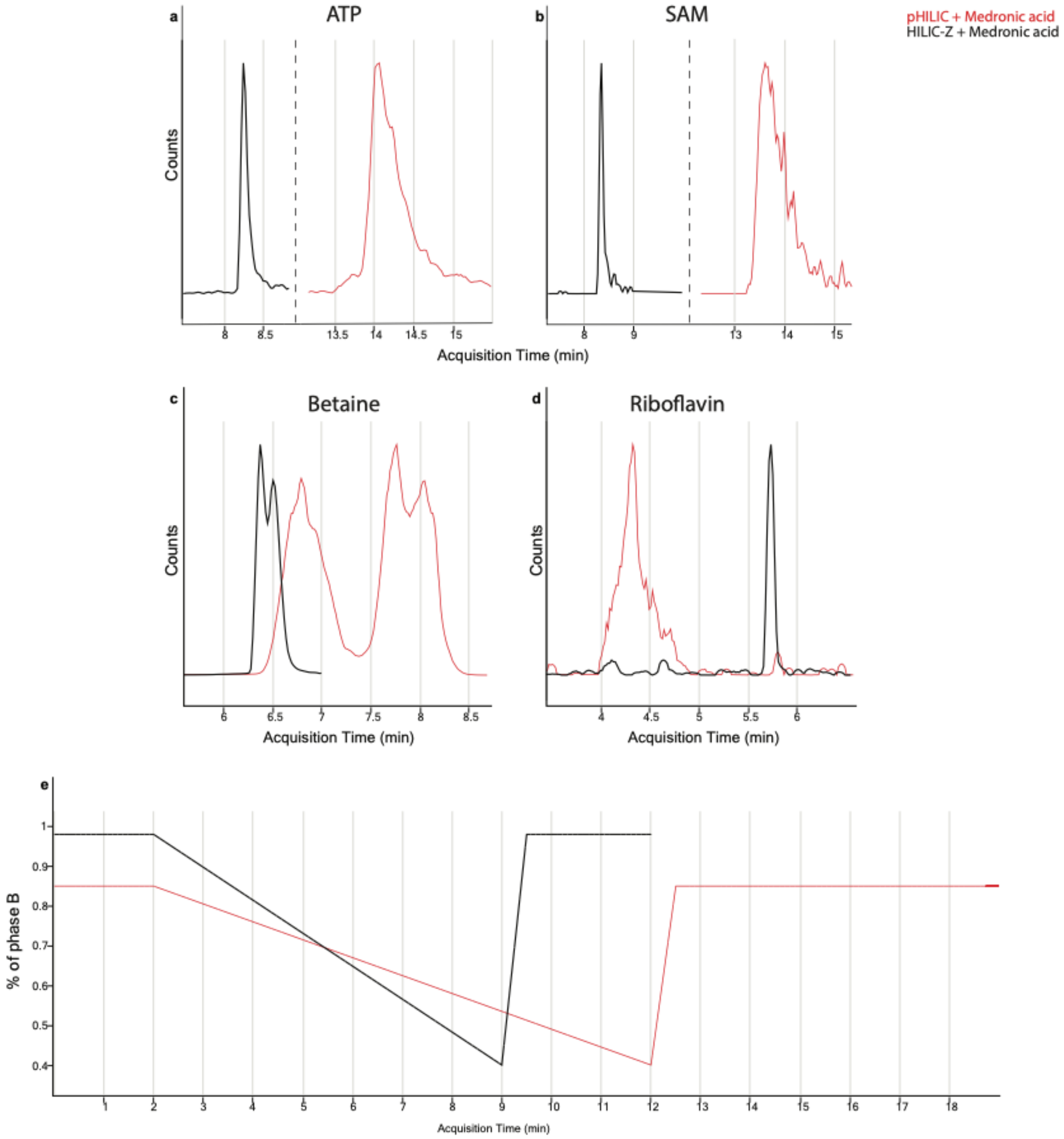


**Supplementary figure 2.** Zwitterionic column comparison.

(a-d) Multiple reaction monitoring (MRM) extracted chromatograms of ATP, SAM, betaine and riboflavin compared using two different zwitterionic columns. (e) Chromatographic elution profiles showing the percentage of B (organic) phase. pHILIC: SeQuant® ZIC-pHILIC column (Merck); HILIC-Z: InfinityLab Poroshell 120 HILIC-Z (Agilent).

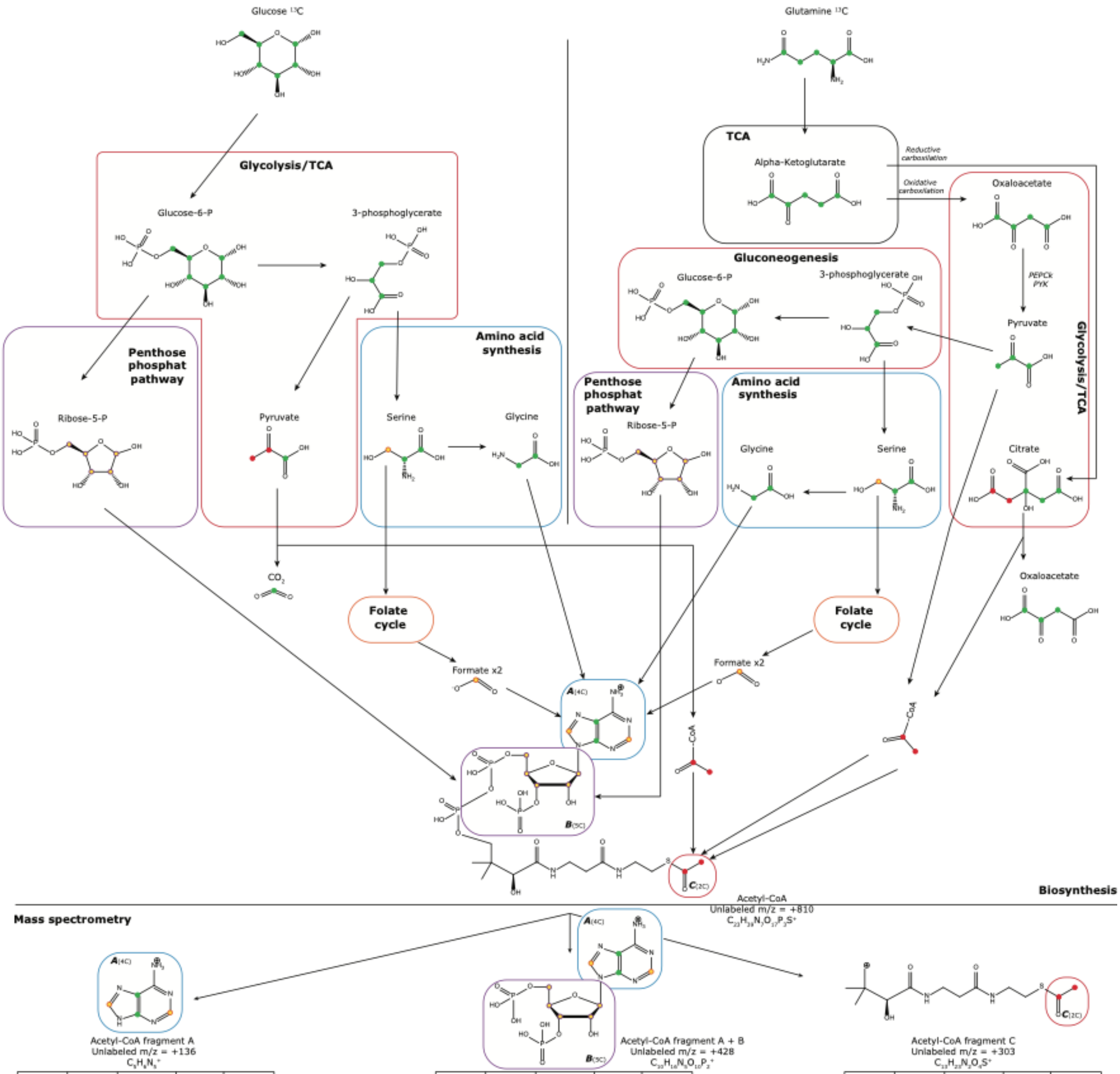


Acetyl-CoA fragment A
Unlabeled m/z = +136
$C_5H_6N_5^+$

| Labeled moiety | Parent ion m/z | Labelling origin | Transition m/z | Monitored ¹³C trans |
|---|---|---|---|---|
| Unlabeled | 810 | -- | 136 | -- |
| A | 811 | +1 (For) | 137 | A |
| A | 812 | +2 (Gly or For) | 138 | A |
| C | 812 | +2 (Ac) | 136 | -- |
| A + C | 813 | +3 (For+Ac) | 137 | A |
| A | 813 | +3 (Gly+For) | 139 | A |
| A | 814 | +4 (Gly+2For) | 140 | A |
| A + C | 814 | +4 (Gly or 2For+Ac) | 138 | A |
| B | 815 | +5 (R5P) | 136 | -- |
| A + C | 815 | +5 (Gly+For +Ac) | 139 | A |
| A + B | 816 | +6 (R5P+For) | 137 | A |
| A + C | 816 | +6 (Gly+2For +Ac) | 140 | A |
| A + B | 817 | +7 (R5P+Gly or 2For) | 138 | A |
| B + C | 817 | +7 (R5P+Ac) | 136 | -- |
| A + B | 818 | +8 (R5P+Gly +For) | 139 | A |
| A + B + C | 818 | +8 (R5P+For +Ac) | 137 | A |
| A + B + C | 819 | +9 (R5P+Gly or 2For+Ac) | 138 | A |
| A + B | 819 | +9 (R5P+Gly +2For) | 140 | A |
| A + B + C | 820 | +10 (R5P+Gly +For+Ac) | 139 | A |
| A + B + C | 821 | +11 (R5P+Gly +2For+Ac) | 140 | A |

For=Formate; Gly=Glycine; Ac=acetyl-CoA; R5P=Ribose-5'-phosphate

Acetyl-CoA fragment A + B
Unlabeled m/z = +428
$C_{10}H_{14}N_5O_{10}P_2^+$

| Labeled moiety | Parent ion m/z | Labelling origin | Transition m/z | Monitored ¹³C trans |
|---|---|---|---|---|
| Unlabeled | 810 | -- | 428 | -- |
| A | 811 | +1 (For) | 429 | A |
| A | 812 | +2 (Gly or For) | 430 | A |
| C | 812 | +2 (Ac) | 428 | -- |
| A + C | 813 | +3 (For+Ac) | 429 | A |
| A | 813 | +3 (Gly+For) | 431 | A |
| A | 814 | +4 (Gly+2For) | 432 | A |
| A + C | 814 | +4 (Gly or 2For+Ac) | 430 | A |
| B | 815 | +5 (R5P) | 433 | B |
| A + C | 815 | +5 (Gly+For +Ac) | 431 | A |
| A + B | 816 | +6 (R5P+For) | 434 | A B |
| A + C | 816 | +6 (Gly+2For +Ac) | 432 | A |
| A + B | 817 | +7 (R5P+Gly or 2For) | 435 | A B |
| B + C | 817 | +7 (R5P+Ac) | 433 | B |
| A + B | 818 | +8 (R5P+Gly +For) | 436 | A B |
| A + B + C | 818 | +8 (R5P+For +Ac) | 434 | A B |
| A + B + C | 819 | +9 (R5P+Gly or 2For+Ac) | 435 | A B |
| A + B | 819 | +9 (R5P+Gly +2For) | 437 | A B |
| A + B + C | 820 | +10 (R5P+Gly +For+Ac) | 436 | A B |
| A + B + C | 821 | +11 (R5P+Gly +2For+Ac) | 437 | A B |

Acetyl-CoA fragment C
Unlabeled m/z = +303
$C_{11}H_{23}N_2O_4S^+$

| Labeled moiety | Parent ion m/z | Labelling origin | Transition m/z | Monitored ¹³C trans |
|---|---|---|---|---|
| Unlabeled | 810 | -- | 303 | -- |
| A | 811 | +1 (For) | 303 | -- |
| A | 812 | +2 (Gly or For) | 303 | -- |
| C | 812 | +2 (Ac) | 305 | C |
| A + C | 813 | +3 (For+Ac) | 305 | C |
| A | 813 | +3 (Gly+For) | 303 | -- |
| A | 814 | +4 (Gly+2For) | 303 | -- |
| A + C | 814 | +4 (Gly or 2For+Ac) | 305 | C |
| B | 815 | +5 (R5P) | 303 | -- |
| A + C | 815 | +5 (Gly+For +Ac) | 305 | C |
| A + B | 816 | +6 (R5P+For) | 303 | -- |
| A + C | 816 | +6 (Gly+2For +Ac) | 305 | C |
| A + B | 817 | +7 (R5P+Gly or 2For) | 303 | -- |
| B + C | 817 | +7 (R5P+Ac) | 305 | C |
| A + B | 818 | +8 (R5P+Gly +For) | 303 | -- |
| A + B + C | 818 | +8 (R5P+For +Ac) | 305 | C |
| A + B + C | 819 | +9 (R5P+Gly or 2For+Ac) | 305 | C |
| A + B | 819 | +9 (R5P+Gly +2For) | 303 | -- |
| A + B + C | 820 | +10 (R5P+Gly +For+Ac) | 305 | C |
| A + B + C | 821 | +11 (R5P+Gly +2For+Ac) | 305 | C |

**Supplementary figure 3.** Acetyl-coenzyme A (acetyl-CoA) moieties biosynthesis, fragmentation patterns and calculated isotopomers from labeled glucose or glutamine.

Biosynthesis and pathways followed by labeled carbon atoms from glucose or glutamine to acetyl-CoA (top). Most intense fragments in the mass spectrometer instrument and calculated isotopomers table (bottom).

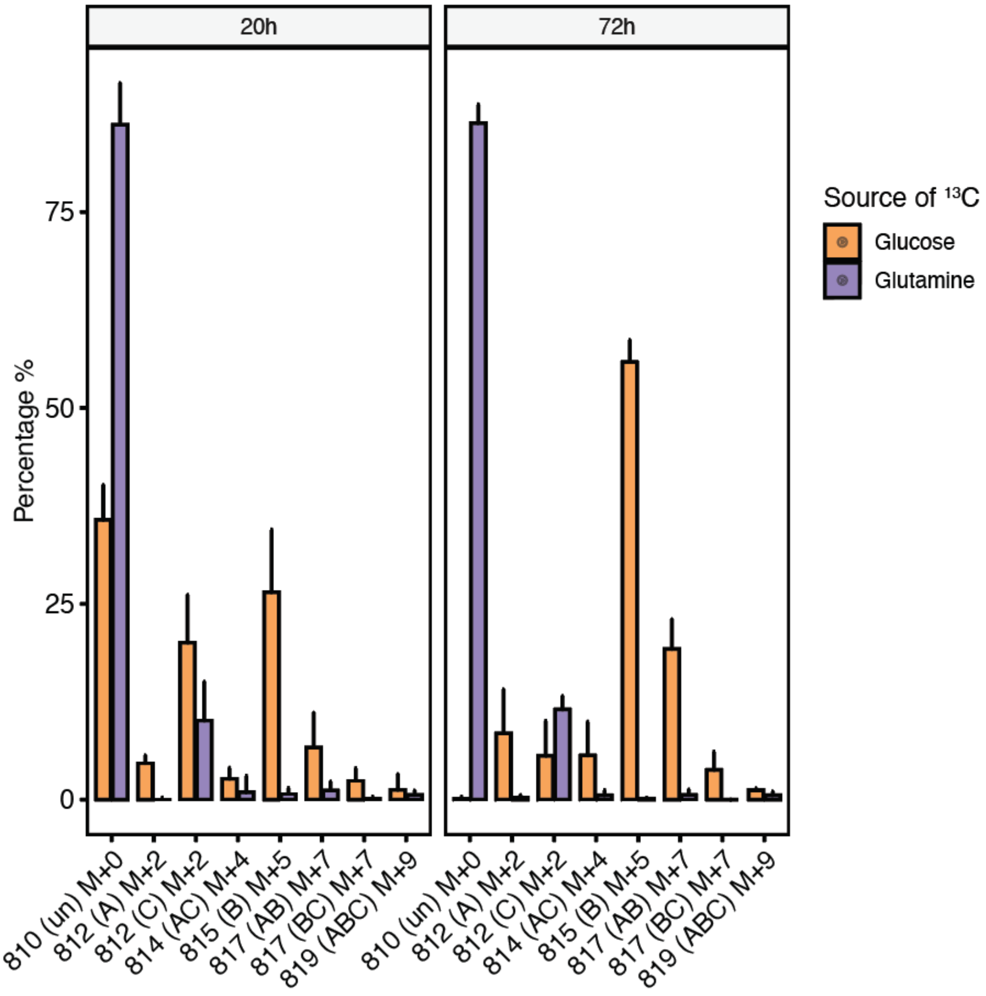


**Supplementary figure 4.** Acetyl-coenzyme A (acetyl-CoA) isotopomers distribution after [U-13C]glucose and [U-13C]glutamine incubation.

Barplots representing isotopomers abundance for acetyl-CoA in an isotope tracing experiment. U-87 MG cells isolated from malignant gliomas were incubated for 20h and 72h in labeled [U-$^{13}C$]glucose or [U-$^{13}C$]glutamine. The x axis shows the m/z of the parent ion, followed by the moiety being labeled between parentheses, followed by the number of extra carbons/neutrons incorporated to the non-labeled compound (M).

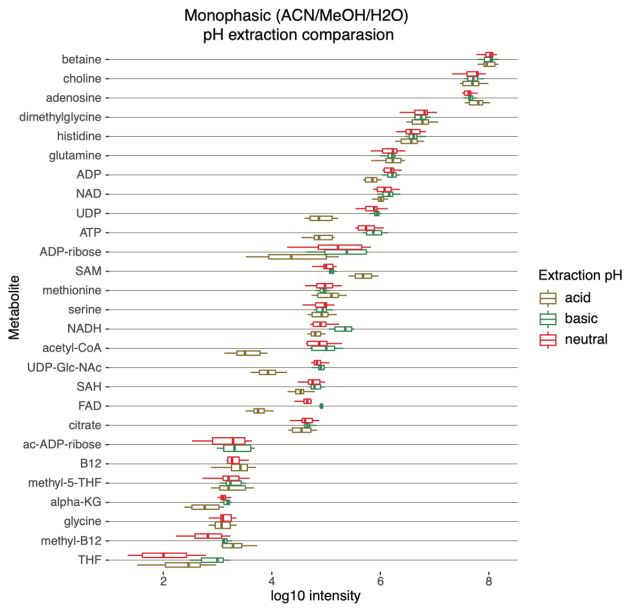


**Supplementary figure 5.** Influence of pH on metabolite extraction.

Liver log10 intensities of metabolites extracted with the acetonitrile:methanol:water (4:4:2, v:v:v) at pH=3.2 (acid), pH=6.6 (neutral) and pH=9.1 (basic) adjusted extraction solutions.

### Supplementary text

**Supplementary text 1**. Rational on how to calculate isotopomer MRM transitions from biological knowledge and non labeled mass spectrum standards

A previous biological knowledge about the origin of the moieties forming the two metabolites tested in the $^{13}C$ labeled flux experiment was required. The two metabolites (SAM and acetyl-CoA) have a ribose moiety where all 5 carbons are susceptible to become labeled when coming from the pentose phosphate pathway, in both, labeled glucose and glutamine. The acetyl group from acetyl-CoA can be labeled from the oxidative carboxylation via oxaloacetate and pyruvate when glucose or glutamine are labeled, but only can be labeled from the reductive carboxylation pathway via alpha-ketoglutarate and citrate, when glutamine was labeled but not glucose. Carbon 4 and 5 in the purine nucleotide adenine present in SAM and acetyl-CoA, can be labeled via glycolysis (from labeled glucose) or gluconeogenesis (from labeled glutamine) through the amino acid synthesis pathway. In SAM, the carbon atom in the methyl group covalently bound to the sulfur atom can be labeled. Only if, the $^{13}C$ is originated from the amino acid serine in the oxidative carboxylation of glutamine or glycolysis from glucose, and through the amino acid synthesis pathway first, and one carbon metabolism later.

Suppose that the most intense fragment of acetyl-CoA for your instrument setup is the $m/z_2$=303, with two possible labeled carbon atoms (Supplementary figure 4). These two carbons are originated from glucose: via glycolysis and through pyruvate; or from glutamine: via oxidative carboxylation through the oxaloacetate-pyruvate pathway, or via reductive carboxylation through citrate, both converting glutamine to alpha-ketoglutaric acid initially. Supposing that only these two labeled carbons were incorporated in the whole acetyl-CoA molecule, we would need to add +2 to the precursor ion $m/z_0$=810 becoming $m/z_{0*}$=812 and the fragment $m/z_2$=303 becoming $m/z_{2*}$=305. But there are at least three moieties in acetyl-CoA that can incorporate labeled carbon atoms from glucose or glutamine (Supplementary figure 3). We take the assumption that all possibly labeled carbon atoms are actually labeled and that there are no intermediate or half labeled moieties. To calculate all labeled possibilities, we assume that fully labeled moieties are independent of each other. We can have labeling in all carbons of all moieties, or not having any labeling in any moiety and all the combinations in between (Table in supplementary figure 3).

Another important step to accurately generate trustable isotope labeling data is to subtract the natural isotopic distribution of previous isotopomers or parent ion in each of the following isotopomers. To measure a more accurate abundance of a parent ion called M+0, and its isotopomers named M+1, M+2...M+n in a glucose carbon $^{13}C$ labeling experiment, the natural isotopic distribution of M+0 need to be taken into account when measuring M+1, M+2..., because will influence the abundance of the ions M+1, M+2...M+n of the same molecule. The natural isotopic distribution of the M+1 (with the influence of the isotope distribution of M+0 already subtracted) will also influence the abundance of the ions M+2, M+3...M+n, and the same for M+2 natural isotopic distribution influencing M+3...M+n and so on. To correct for the contribution of naturally occurring isotope distributions among the isotopomers that are monitored, the natural isotope distribution of the previous parent or isotopologue need to be subtracted. To do so, the theoretical natural isotope distribution for each of the m/z monitored (parent ion or previous isotopologue) were obtained using the enviPat web at https://www.envipat.eawag.ch/. From which the highest pattern profile was obtained for each parent (or isotopologue) using the molecular formula and taking into account the number of carbon $^{13}C$ atoms present. The influence isotopic threshold was set to $1x10^{-10}$ in order to account for most of the natural distribution of $^{13}C$ atoms. Once this matrix with contributions in the isotopic distributions of all monitored m/z values for the same formula were obtained, the following operations were performed up until the last isotopologue species:

$(M+0)_r = (M+0)_m$ => the M+0 has no natural isotopic distribution influence from a predecessor isotopologue with the same exact chemical formula.

$(M+1)_r = (M+1)_m - ((M+0)_m * (M+0)_1 / 100)$

$(M+2)_i = (M+2)_m - ((M+0)_m * (M+0)_2 / 100)$

$(M+2)_r = (M+2)_i - ((M+1)_r * (M+1)_1 / 100)$

$(M+3)_i = (M+3)_m - ((M+0)_m * (M+0)_3 / 100)$

$(M+3)_{ii} = (M+3)_i - ((M+1)_r * (M+1)_2 / 100)$

$(M+3)_r = (M+3)_{ii} - ((M+2)_r * (M+1)_1 / 100)$

where M + (isotopologue number)$_m$ represents the measured abundance for the M + (isotopologue number), M + (isotopologue number)$_{i \text{ or } ii \text{ or } iii...}$represents the intermediate abundances for the M + (isotopologue number), M + (isotopologue number)$_r$ represents the real abundances for that particular M + (isotopologue number) after subtracting the corresponding influences of predecessor isotopomers or parent ion, and where M + (isotopologue number)$_{1 \text{ or } 2 \text{ or } 3...}$represents the theoretical abundances of the isotopomers for that particular specie calculated with enviPat expressed in percentage.

### Supplementary tables

**Supplementary table 1.** Epigenetically relevant metabolites complete information.

Metabolite name, chemical formula, monoisotopic mass, 1$^{st}$ transition (collision energy), 2$^{nd}$ transition (collision energy), extraction with best peak shape, polarity detected, chromatography used, origin of the metabolite and highest contribution for the host, reported epigenetic function/s and references.

| Metabolite | Chemical formula | Monoisotopic mass (g/mol) | Parent m/z | m/z 1st transition (CE) | m/z 2nd transition (CE) | Type of extraction with better peak shape | Polarity mode of detection | Metabolomics analytical setup | Origin (Highest contribution for the host) | Epigenetic Function (at least one) | Reference |
|---|---|---|---|---|---|---|---|---|---|---|---|
| **Formic acid** | $CH_2O_2$ | 46.005479 | 46 | 45() | 29() | Biphasic | Pos | GC-QqQ | Microbiome | Methyl donor | [35] |
| **Acetic acid** | $C_2H_4O_2$ | 60.021129 | 60 | 45() | 43() | Biphasic | Pos | GC-QqQ | Microbiome | Acetyl donor | [41] |
| **Propionic acid** | $C_3H_6O_2$ | 74.036779 | 74 | 73() | 55() | Biphasic | Pos | GC-QqQ | Microbiome | Acetyl donor and HDAC3 inhibitor | [15,42,43] |
| **Butyric acid** | $C_4H_8O_2$ | 88.052429 | 88 | 60() | 73() | Biphasic | Pos | GC-QqQ | Microbiome | Acetyl donor and HDACs inhibitor | [29,44–47] |
| **Glycine** | $C_2H_5NO_2$ | 75.032028 | 76 | 30 (8) | 48 (4) | Monophasic | Pos | LC-QqQ | Host metabolism | Methyl donor | [48] |
| **Choline** | $C_5H_{14}NO^+$ | 104.107539 | 104 | 60 (20) | 45 (15) | Both | Pos | LC-QqQ | Diet | Methyl donor | [49] |
| **N, N-Dimethylglycine** | $C_4H_9NO_2$ | 103.063329 | 104 | 58 (12) | 42 (35) | Both | Pos | LC-QqQ | Host metabolism | Methylation transfer product | [50] |
| **Serine** | $C_3H_7NO_3$ | 105.042593 | 106 | 60 (12) | 42 (20) | Both | Pos | LC-QqQ | Host metabolism | Methyl donor | [48] |
| **Fumaric acid** | $C_4H_4O_4$ | 116.010959 | -115 | -71 (4) | -27 (8) | Both | Neg | LC-QqQ | Host metabolism | Inhibitor of TET and JmjC demethylation enzymes | [51] |
| **Succinic acid** | $C_4H_6O_4$ | 118.02609 | -117 | -73 (8) | -99 (8) | Both | Neg | LC-QqQ | Host and microbiome | Inhibitor of TET and JmjC demethylation enzymes | [24,52,53] |
| **Betaine** | $C_5H_{11}NO_2$ | 117.078979 | 118 | 58 (28) | 59 (16) | Both | Pos | LC-QqQ | Diet | Methyl donor | [54] |
| **Homocysteine** | $C_4H_9NO_2S$ | 135.0354 | 136 | 90 (8) | 56 (16) | none | Pos | LC-QqQ | Host metabolism | Methyl acceptor | [50] |
| **Alpha-Ketoglutaric acid** | $C_5H_4O_5^{-2}$ | 144.005873 | -145 | -101 (4) | -57 (8) | Both | Neg | LC-QqQ | Host metabolism | JmjC co-substrate | [9,10,55] |
| **Glutamine** | $C_5H_{10}N_2O_3$ | 146.069142 | 147 | 84 (20) | 130 (8) | Both | Pos | LC-QqQ | Host metabolism and diet | Alpha-Ketoglutaric acid precursor | [56] |
| **Methionine** | $C_5H_{11}NO_2S$ | 149.05105 | 150 | 56 (12) | 104 (6) | Both | Pos | LC-QqQ | Diet | Methyl donor | [28] |
| **Histidine** | $C_6H_9N_3O_2$ | 155.069477 | 156 | 110 (12) | 83 (28) | Both | Pos | LC-QqQ | Diet | THF precursor | [11] |
| **Glucose** | $C_6H_{12}O_6$ | 180.063388 | 181 | 99 (12) | 140 (4) | Biphasic | Pos | LC-QqQ | Diet and host metabolism | TET enzyme modulator | [57] |

| | | | | | | | | | | | |
|---|---|---|---|---|---|---|---|---|---|---|---|
| **Citrate** | $C_6H_5O_7^{-3}$ | 189.003527 | -191 | -67 (28) | -57 (20) | Both | Neg | LC-QqQ | Host metabolism | Acetyl donor | [58] |
| **Pantothenic acid** **(Vitamin B5)** | $C_9H_{17}NO_5$ | 219.110673 | 220 | 90 (12) | 202 (12) | Both | Pos | LC-QqQ | Diet and microbiome | Acetyl donor | [59] |
| **N-acetylglucosamine** | $C_8H_{15}NO_6$ | 221.089937 | 222 | 138 (12) | 204 (4) | Biphasic | Pos | LC-QqQ | Diet? | PRC2 substrate and GlcNAc unit | [60,61] |
| **Adenosine** | $C_{10}H_{13}N_5O_4$ | 267.096754 | 268 | 136 (16) | 119 (48) | Both | Pos | LC-QqQ | Host metabolism and microbiome? | DNMT inhibitor | [62] |
| **Adenosine 5'-monophosphate (AMP)** | $C_{10}H_{14}N_5O_7P$ | 347.063085 | 348 | 136 (15) | 97 (30) | Both | Pos | LC-QqQ | Host metabolism | Phosphor donor | [63] |
| **Riboflavin** **(Vitamin B2)** | $C_{17}H_{20}N_4O_6$ | 376.138284 | 377 | 243 (20) | 172 (35) | Both | Pos | LC-QqQ | Diet and microbiome | FAD precursor and LSD | [27,59] |
| **S-adenosylhomocysteine (SAH)** | $C_{14}H_{20}N_6O_5S$ | 384.121589 | 385 | 136 (24) | 250 (4) | Both | Pos | LC-QqQ | Host metabolism and microbiome? | Methylation transfer product and methyltransferase s inhibitor | [27] |
| **S-adenosylmethionine (SAM)** | $C_{15}H_{23}N_6O_5S^+$ | 399.145064 | 399 | 250 (12) | 97 (32) | Monophasic | Pos | LC-QqQ | Host metabolism and microbiome? | Methyl donor and co-factor | [27] |
| **Uridine 5'-diphosphate (UDP)** | $C_9H_{14}N_2O_{12}P_2$ | 404.002198 | 405 | 97 (24) | 113 (36) | Monophasic | Pos | LC-QqQ | Host metabolism | GlcNAc carrier | [64] |
| **Adenosine 5'-diphosphate (ADP)** | $C_{10}H_{15}N_5O_{10}P_2$ | 427.029416 | 428 | 136 (32) | 348 (16) | Monophasic | Pos | LC-QqQ | Host metabolism | Ribosylation carrier | [65] |
| **Dihydrofolic acid (DHF)** | $C_{19}H_{21}N_7O_6$ | 443.155331 | -442 | -176 (28) | -265 (16) | Monophasic | Neg | LC-QqQ | Microbiome and host metabolism | Methyl donor | [11] |
| **Folic acid (Vitamin B9)** | $C_{19}H_{19}N_7O_6$ | 441.139681 | 442 | 295 (12) | 120 (40) | Biphasic | Pos | LC-QqQ | Microbiome and host metabolism | Methyl donor | [11,59,66] |
| **Tetrahydrofolic acid (THF)** | $C_{19}H_{23}N_7O_6$ | 445.170981 | 446 | 299 (20) | 120 (44) | Monophasic | Pos | LC-QqQ | Host metabolism and microbiome | Methyl donor | [11] |

| | | | | | | | | | | | |
|---|---|---|---|---|---|---|---|---|---|---|---|
| **N5-Methyl-tetrahydrofolic acid** | $C_{20}H_{25}N_7O_6$ | 459.186632 | 460 | 313 (24) | 180 (44) | Monophasic | Pos | LC-QqQ | Host metabolism and microbiome | Methyl donor | [11] |
| **Adenosine 5'-triphosphate (ATP)** | $C_{10}H_{16}N_5O_{13}P_3$ | 506.995747 | 508 | 136 (44) | 97 (40) | Both | Pos | LC-QqQ | Host metabolism | Phosphate donor and co-factor | [65] |
| **ADP Ribose (ADPR)** | $C_{15}H_{23}N_5O_{14}P_2$ | 559.071674 | 560 | 136 (36) | 348 (16) | Monophasic | Pos | LC-QqQ | Host metabolism | Ribose donor and acetyl group acceptor | [65] |
| **O-Acetyl-ADP-Ribose** | $C_{17}H_{25}N_5O_{15}P_2$ | 601.082239 | 602 | 136 (36) | 348 (20) | Both | Pos | LC-QqQ | Host metabolism | Acetyl donor | [65] |
| **UDP-N-acetylglucosamine** | $C_{17}H_{27}N_3O_{17}P_2$ | 607.08157 | 608 | 204 (8) | 138 (48) | Monophasic | Pos | LC-QqQ | Host metabolism | GlcNAc donor | [64] |
| **Nicotinamide adenine dinucleotide (NAD)** | $C_{21}H_{28}N_7O_{14}P_2^+$ | 664.116948 | 664 | 136 (60) | 428 (24) | Monophasic | Pos | LC-QqQ | Host metabolism | Deacetylases and Sirtuins cofactor | [67,68] |
| **NADH** | $C_{21}H_{29}N_7O_{14}P_2$ | 665.124773 | 666 | 136 (40) | 137 (52) | Monophasic | Pos | LC-QqQ | Host metabolism | Reduced NAD form | [67,68] |
| **Coenzyme A (CoA)** | $C_{21}H_{36}N_7O_{16}P_3S$ | 767.11521 | 768 | 261 (30) | 428 (25) | Both | Pos | LC-QqQ | | Acetyl group carrier, pantothenic acid precursor and coenzyme | [59] |
| **Flavin adenine dinucleotide (FAD)** | $C_{27}H_{33}N_9O_{15}P_2$ | 785.157135 | 786 | 348 (24) | 136 (48) | Monophasic | Pos | LC-QqQ | Diet | LSD demethylases cofactor | [27] |
| **Acetyl-Coenzyme A (ac-CoA)** | $C_{23}H_{38}N_7O_{17}P_3S$ | 809.125775 | 810 | 303 (36) | 136 (60) | Monophasic | Pos | LC-QqQ | Host metabolism | Acetyl donor and acetyltransferase cofactor | [13,67] |
| **Malonyl-CoA** | $C_{24}H_{38}N_7O_{19}P_3S$ | 853.115604 | 854 | 303 (32) | 347 (48) | Biphasic | Pos | LC-QqQ | Host metabolism | Malonyl donor | [69,70] |
| **Cyanocobalamin** (Vitamin B12) | $C_{63}H_{88}CoN_{14}O_{14}P$ | 1355.57522 | 678 | 147 (44) | 912 (40) | Biphasic | Pos | LC-QqQ | Diet and microbiome | Methyltransferase cofactor | [26] |

**Supplementary table 2.** SAM and acetyl-CoA isotopomer moieties further details.

Parent ions, transitions and collision energies for labeled SAM and Acetyl-CoA different moieties.

| Metabolite | Labeled moieties | Parent m/z | Transition m/z & (collision energy) |
|---|---|---|---|
| SAM | NA | 399 | 97 (32) |
| | NA | 399 | 136 (24) |
| | NA | 399 | 250 (12) |
| | NA | 399 | 298 (4) |
| | A | 400 | 97 (32) |
| | C | 400 | 97 (32) |
| | A | 401 | 97 (32) |
| | A + C | 401 | 97 (32) |
| | A | 402 | 97 (32) |
| | A + C | 402 | 97 (32) |
| | A | 403 | 97 (32) |
| | A + C | 403 | 97 (32) |
| | B | 404 | 102 (32) |
| | A + C | 404 | 97 (32) |
| | A + B | 405 | 102 (32) |
| | B + C | 405 | 102 (32) |
| | A + B | 406 | 102 (32) |
| | A + B + C | 406 | 102 (32) |
| | A + B + C | 407 | 102 (32) |
| | A + B | 407 | 102 (32) |
| | A + B | 408 | 102 (32) |

| | | | |
|---|---|---|---|
| | A + B + C | 408 | 102 (32) |
| | A + B + C | 409 | 102 (32) |
| | A | 400 | 137 (24) |
| | C | 400 | 136 (24)) |
| | A | 401 | 138 (24) |
| | A + C | 401 | 137 (24) |
| | A | 402 | 139 (24) |
| | A + C | 402 | 138 (24) |
| | A | 403 | 140 (24) |
| | A + C | 403 | 139 (24) |
| | B | 404 | 136 (24) |
| | A + C | 404 | 140 (24) |
| | A + B | 405 | 137 (24) |
| | B + C | 405 | 136 (24) |
| | A + B | 406 | 138 (24) |
| | A + B + C | 406 | 137 (24) |
| | A + B + C | 407 | 138 (24) |
| | A + B | 407 | 139 (24) |
| | A + B | 408 | 140 (24) |
| | A + B + C | 408 | 139 (24) |
| | A + B + C | 409 | 140 (24) |
| | A | 400 | 251 (12) |
| | C | 400 | 250 (12) |
| | A | 401 | 252 (12) |
| | A + C | 401 | 251 (12) |
| | A | 402 | 253 (12) |

| | | | |
|---|---|---|---|
| | A + C | 402 | 252 (12) |
| | A | 403 | 254 (12) |
| | A + C | 403 | 253 (12) |
| | B | 404 | 255 (12) |
| | A + C | 404 | 254 (12) |
| | A + B | 405 | 256 (12) |
| | B + C | 405 | 255 (12) |
| | A + B | 406 | 257 (12) |
| | A + B + C | 406 | 256 (12) |
| | A + B + C | 407 | 257 (12) |
| | A + B | 407 | 258 (12) |
| | A + B | 408 | 259 (12) |
| | A + B + C | 408 | 258 (12) |
| | A + B + C | 409 | 259 (12) |
| | A | 400 | 299 (4) |
| | C | 400 | 299 (4) |
| | A | 401 | 300 (4) |
| | A + C | 401 | 300 (4) |
| | A | 402 | 301 (4) |
| | A + C | 402 | 301 (4) |
| | A | 403 | 302 (4) |
| | A + C | 403 | 302 (4) |
| | B | 404 | 303 (4) |
| | A + C | 404 | 303 (4) |
| | A + B | 405 | 304 (4) |
| | B + C | 405 | 304 (4) |

| | | | |
|---|---|---|---|
| | A + B | 406 | 305 (4) |
| | A + B + C | 406 | 305 (4) |
| | A + B + C | 407 | 306 (4) |
| | A + B | 407 | 306 (4) |
| | A + B | 408 | 307 (4) |
| | A + B + C | 408 | 307 (4) |
| | A + B + C | 409 | 308 (4) |
| Acetyl-CoA | NA | 810 | 136 (60) |
| | NA | 810 | 303 (36) |
| | NA | 810 | 428 (25) |
| | A | 811 | 137 (60) |
| | A | 812 | 138 (60) |
| | C | 812 | 136 (60) |
| | A + C | 813 | 137 (60) |
| | A | 813 | 139 (60) |
| | A | 814 | 140 (60) |
| | A + C | 814 | 138 (60) |
| | B | 815 | 136 (60) |
| | A + C | 815 | 139 (60) |
| | A + B | 816 | 137 (60) |
| | A + C | 816 | 140 (60) |
| | A + B | 817 | 138 (60) |
| | B + C | 817 | 136 (60) |
| | A + B | 818 | 139 (60) |
| | A + B + C | 818 | 137 (60) |
| | A + B + C | 819 | 138 (60) |

| | | | |
|---|---|---|---|
| | A + B | 819 | 140 (60) |
| | A + B + C | 820 | 139 (60) |
| | A + B + C | 821 | 140 (60) |
| | A | 811 | 303 (36) |
| | A | 812 | 303 (36) |
| | C | 812 | 305 (36) |
| | A + C | 813 | 305 (36) |
| | A | 813 | 303 (36) |
| | A | 814 | 303 (36) |
| | A + C | 814 | 305 (36) |
| | B | 815 | 303 (36) |
| | A + C | 815 | 305 (36) |
| | A + B | 816 | 303 (36) |
| | A + C | 816 | 305 (36) |
| | A + B | 817 | 303 (36) |
| | B + C | 817 | 305 (36) |
| | A + B | 818 | 303 (36) |
| | A + B + C | 818 | 305 (36) |
| | A + B + C | 819 | 305 (36) |
| | A + B | 819 | 303 (36) |
| | A + B + C | 820 | 305 (36) |
| | A + B + C | 821 | 305 (36) |
| | A | 811 | 429 (25) |
| | A | 812 | 430 (25) |
| | C | 812 | 428 (25) |
| | A + C | 813 | 429 (25) |

| | | |
|---|---|---|
| A | 813 | 431 (25) |
| A | 814 | 432 (25) |
| A + C | 814 | 430 (25) |
| B | 815 | 433 (25) |
| A + C | 815 | 431 (25) |
| A + B | 816 | 434 (25) |
| A + C | 816 | 432 (25) |
| A + B | 817 | 435 (25) |
| B + C | 817 | 433 (25) |
| A + B | 818 | 436 (25) |
| A + B + C | 818 | 434 (25) |
| A + B + C | 819 | 435 (25) |
| A + B | 819 | 437 (25) |
| A + B + C | 820 | 436 (25) |
| A + B + C | 821 | 437 (25) |

**Supplementary table 3.** [U-$^{13}$C]glucose and [U-$^{13}$C]glutamine labeled medium used for SAM and acetyl-CoA isotope trace experiments.

**[U-$^{13}$C]glucose Complete medium for acetyl-CoA**

| | |
|---|---|
| Nacalai-Tesque 09848 | 500 mL |
| $^{13}$C-Glucose 1 g/l | |
| Sodium Pyruvate 0.11 g/l = 1 mM | 5 mL from 100 mM stock |
| FBS | 50 mL |

**[U-$^{13}$C]glutamine Complete medium for acetyl-CoA**

| | |
|---|---|
| Sigma M5650 | 500 mL |
| $^{13}$C-Glutamine 0.292 g/l = 2 mM (stock 5 mg) | |
| Sodium Pyruvate 0.11 g/l = 1 mM | 5 mL from 100 mM stock |
| FBS | 50 mL |

(20) Liu, X.; Sadhukhan, S.; Sun, S.; Wagner, G. R.; Hirschey, M. D.; Qi, L.; Lin, H.; Locasale, J. W. High-Resolution Metabolomics with Acyl-CoA Profiling Reveals Widespread Remodeling in

Response to Diet. *Mol. Cell. Proteomics* **2015**, *14* (6), 1489–1500.
(21) Wright, R. H. G.; Lioutas, A.; Le Dily, F.; Soronellas, D.; Pohl, A.; Bonet, J.; Nacht, A. S.; Samino, S.; Font-Mateu, J.; Vicent, G. P.; Wierer, M.; Trabado, M. A.; Schelhorn, C.; Carolis, C.; Macias, M. J.; Yanes, O.; Oliva, B.; Beato, M. ADP-Ribose-Derived Nuclear ATP Synthesis by NUDIX5 Is Required for Chromatin Remodeling. *Science* **2016**, *352* (6290), 1221–1225.
(22) Schittmayer, M.; Birner-Gruenberger, R.; Zamboni, N. Quantification of Cellular Folate Species by LC-MS after Stabilization by Derivatization. *Anal. Chem.* **2018**, *90* (12), 7349–7356.
(23) Lotti, C.; Rubert, J.; Fava, F.; Tuohy, K.; Mattivi, F.; Vrhovsek, U. Development of a Fast and Cost-Effective Gas Chromatography-Mass Spectrometry Method for the Quantification of Short-Chain and Medium-Chain Fatty Acids in Human Biofluids. *Anal. Bioanal. Chem.* **2017**, *409* (23), 5555–5567.
(24) Yang, M.; Pollard, P. J. Succinate: A New Epigenetic Hacker. *Cancer Cell* **2013**, *23* (6), 709–711.
(25) Shuvalov, O.; Petukhov, A.; Daks, A.; Fedorova, O.; Vasileva, E.; Barlev, N. A. One-Carbon Metabolism and Nucleotide Biosynthesis as Attractive Targets for Anticancer Therapy. *Oncotarget* **2017**, *8* (14), 23955–23977.
(26) Rush, E. C.; Katre, P.; Yajnik, C. S. Vitamin B12: One Carbon Metabolism, Fetal Growth and Programming for Chronic Disease. *Eur. J. Clin. Nutr.* **2014**, *68* (1), 2–7.
(27) Kaelin, W. G., Jr; McKnight, S. L. Influence of Metabolism on Epigenetics and Disease. *Cell* **2013**, *153* (1), 56–69.
(28) Maddocks, O. D. K.; Labuschagne, C. F.; Adams, P. D.; Vousden, K. H. Serine Metabolism Supports the Methionine Cycle and DNA/RNA Methylation through De Novo ATP Synthesis in Cancer Cells. *Mol. Cell* **2016**, *61* (2), 210–221.
(29) Chang, P. V.; Hao, L.; Offermanns, S.; Medzhitov, R. The Microbial Metabolite Butyrate Regulates Intestinal Macrophage Function via Histone Deacetylase Inhibition. *Proc. Natl. Acad. Sci. U. S. A.* **2014**, *111* (6), 2247–2252.
(30) Miro-Blanch, J.; Yanes, O. Epigenetic Regulation at the Interplay Between Gut Microbiota and Host Metabolism. *Front. Genet.* **2019**, *10*, 638.
(31) Zeng, M.; Cao, H. Fast Quantification of Short Chain Fatty Acids and Ketone Bodies by Liquid Chromatography-Tandem Mass Spectrometry after Facile Derivatization Coupled with Liquid-Liquid Extraction. *J. Chromatogr. B Analyt. Technol. Biomed. Life Sci.* **2018**, *1083*, 137–145.
(32) Lamarre, S. G.; MacMillan, L.; Morrow, G. P.; Randell, E.; Pongnopparat, T.; Brosnan, M. E.; Brosnan, J. T. An Isotope-Dilution, GC-MS Assay for Formate and Its Application to Human and Animal Metabolism. *Amino Acids* **2014**, *46* (8), 1885–1891.
(33) Cummings, J. H.; Pomare, E. W.; Branch, W. J.; Naylor, C. P.; Macfarlane, G. T. Short Chain Fatty Acids in Human Large Intestine, Portal, Hepatic and Venous Blood. *Gut* **1987**, *28* (10), 1221–1227.
(34) Hughes, E. R.; Winter, M. G.; Duerkop, B. A.; Spiga, L.; Furtado de Carvalho, T.; Zhu, W.; Gillis, C. C.; Büttner, L.; Smoot, M. P.; Behrendt, C. L.; Cherry, S.; Santos, R. L.; Hooper, L. V.; Winter, S. E. Microbial Respiration and Formate Oxidation as Metabolic Signatures of Inflammation-Associated Dysbiosis. *Cell Host Microbe* **2017**, *21* (2), 208–219.
(35) Pietzke, M.; Meiser, J.; Vazquez, A. Formate Metabolism in Health and Disease. *Mol Metab* **2020**, *33*, 23–37.
(36) Jang, C.; Chen, L.; Rabinowitz, J. D. Metabolomics and Isotope Tracing. *Cell* **2018**, *173* (4), 822–837.
(37) Kovatcheva, M.; Melendez, E.; Chondronasiou, D.; Pietrocola, F.; Bernad, R.; Caballe, A.; Junza, A.; Capellades, J.; Holguín-Horcajo, A.; Prats, N.; Durand, S.; Rovira, M.; Yanes, O.; Stephan-Otto Attolini, C.; Kroemer, G.; Serrano, M. Vitamin B12 Is a Limiting Factor for Induced Cellular Plasticity and Tissue Repair. *Nature Metabolism* **2023**, 1–20.
(38) Yu, W.; Wang, Z.; Zhang, K.; Chi, Z.; Xu, T.; Jiang, D.; Chen, S.; Li, W.; Yang, X.; Zhang, X.; Wu, Y.; Wang, D. One-Carbon Metabolism Supports S-Adenosylmethionine and Histone Methylation to Drive Inflammatory Macrophages. *Mol. Cell* **2019**, *75* (6), 1147–1160.e5.
(39) Boyle, J. Lehninger Principles of Biochemistry (4th Ed.): Nelson, D., and Cox, M. *Biochemistry and Molecular Biology Education*. 2005, pp 74–75. https://doi.org/10.1002/bmb.2005.494033010419.
(40) Yuan, M.; Breitkopf, S. B.; Yang, X.; Asara, J. M. A Positive/negative Ion-Switching, Targeted Mass

Spectrometry-Based Metabolomics Platform for Bodily Fluids, Cells, and Fresh and Fixed Tissue. *Nat. Protoc.* **2012**, *7* (5), 872–881.
(41) Gao, X.; Lin, S.-H.; Ren, F.; Li, J.-T.; Chen, J.-J.; Yao, C.-B.; Yang, H.-B.; Jiang, S.-X.; Yan, G.-Q.; Wang, D.; Wang, Y.; Liu, Y.; Cai, Z.; Xu, Y.-Y.; Chen, J.; Yu, W.; Yang, P.-Y.; Lei, Q.-Y. Acetate Functions as an Epigenetic Metabolite to Promote Lipid Synthesis under Hypoxia. *Nat. Commun.* **2016**, *7*, 11960.
(42) Lee, E. S.; Song, E. J.; Nam, Y. D. Dysbiosis of Gut Microbiome and Its Impact on Epigenetic Regulation. *J Clin Epigenet.*
(43) Chen, Y.; Sprung, R.; Tang, Y.; Ball, H.; Sangras, B.; Kim, S. C.; Falck, J. R.; Peng, J.; Gu, W.; Zhao, Y. Lysine Propionylation and Butyrylation Are Novel Post-Translational Modifications in Histones. *Mol. Cell. Proteomics* **2007**, *6* (5), 812–819.
(44) Furusawa, Y.; Obata, Y.; Fukuda, S.; Endo, T. A.; Nakato, G.; Takahashi, D.; Nakanishi, Y.; Uetake, C.; Kato, K.; Kato, T.; Takahashi, M.; Fukuda, N. N.; Murakami, S.; Miyauchi, E.; Hino, S.; Atarashi, K.; Onawa, S.; Fujimura, Y.; Lockett, T.; Clarke, J. M.; Topping, D. L.; Tomita, M.; Hori, S.; Ohara, O.; Morita, T.; Koseki, H.; Kikuchi, J.; Honda, K.; Hase, K.; Ohno, H. Commensal Microbe-Derived Butyrate Induces the Differentiation of Colonic Regulatory T Cells. *Nature* **2013**, *504* (7480), 446–450.
(45) Koh, A.; De Vadder, F.; Kovatcheva-Datchary, P.; Bäckhed, F. From Dietary Fiber to Host Physiology: Short-Chain Fatty Acids as Key Bacterial Metabolites. *Cell* **2016**, *165* (6), 1332–1345.
(46) Berni Canani, R.; Di Costanzo, M.; Leone, L. The Epigenetic Effects of Butyrate: Potential Therapeutic Implications for Clinical Practice. *Clin. Epigenetics* **2012**, *4* (1), 4.
(47) Fellows, R.; Denizot, J.; Stellato, C.; Cuomo, A.; Jain, P.; Stoyanova, E.; Balázsi, S.; Hajnády, Z.; Liebert, A.; Kazakevych, J.; Blackburn, H.; Corrêa, R. O.; Fachi, J. L.; Sato, F. T.; Ribeiro, W. R.; Ferreira, C. M.; Perée, H.; Spagnuolo, M.; Mattiuz, R.; Matolcsi, C.; Guedes, J.; Clark, J.; Veldhoen, M.; Bonaldi, T.; Vinolo, M. A. R.; Varga-Weisz, P. Microbiota Derived Short Chain Fatty Acids Promote Histone Crotonylation in the Colon through Histone Deacetylases. *Nat. Commun.* **2018**, *9* (1), 105.
(48) Locasale, J. W. Serine, Glycine and One-Carbon Units: Cancer Metabolism in Full Circle. *Nat. Rev. Cancer* **2013**, *13* (8), 572–583.
(49) Romano, K. A.; Martinez-Del Campo, A.; Kasahara, K.; Chittim, C. L.; Vivas, E. I.; Amador-Noguez, D.; Balskus, E. P.; Rey, F. E. Metabolic, Epigenetic, and Transgenerational Effects of Gut Bacterial Choline Consumption. *Cell Host Microbe* **2017**, *22* (3), 279–290.e7.
(50) Obeid, R. The Metabolic Burden of Methyl Donor Deficiency with Focus on the Betaine Homocysteine Methyltransferase Pathway. *Nutrients* **2013**, *5* (9), 3481–3495.
(51) Sciacovelli, M.; Gonçalves, E.; Johnson, T. I.; Zecchini, V. R.; da Costa, A. S. H.; Gaude, E.; Drubbel, A. V.; Theobald, S. J.; Abbo, S. R.; Tran, M. G. B.; Rajeeve, V.; Cardaci, S.; Foster, S.; Yun, H.; Cutillas, P.; Warren, A.; Gnanapragasam, V.; Gottlieb, E.; Franze, K.; Huntly, B.; Maher, E. R.; Maxwell, P. H.; Saez-Rodriguez, J.; Frezza, C. Fumarate Is an Epigenetic Modifier That Elicits Epithelial-to-Mesenchymal Transition. *Nature* **2016**, *537* (7621), 544–547.
(52) Serena, C.; Ceperuelo-Mallafré, V.; Keiran, N.; Queipo-Ortuño, M. I.; Bernal, R.; Gomez-Huelgas, R.; Urpi-Sarda, M.; Sabater, M.; Pérez-Brocal, V.; Andrés-Lacueva, C.; Moya, A.; Tinahones, F. J.; Fernández-Real, J. M.; Vendrell, J.; Fernández-Veledo, S. Elevated Circulating Levels of Succinate in Human Obesity Are Linked to Specific Gut Microbiota. *ISME J.* **2018**, *12* (7), 1642–1657.
(53) Nag, A.; St John, P. C.; Crowley, M. F.; Bomble, Y. J. Prediction of Reaction Knockouts to Maximize Succinate Production by Actinobacillus Succinogenes. *PLoS One* **2018**, *13* (1), e0189144.
(54) Sun, S.; Li, X.; Ren, A.; Du, M.; Du, H.; Shu, Y.; Zhu, L.; Wang, W. Choline and Betaine Consumption Lowers Cancer Risk: A Meta-Analysis of Epidemiologic Studies. *Sci. Rep.* **2016**, *6*, 35547.
(55) Carey, B. W.; Finley, L. W. S.; Cross, J. R.; Allis, C. D.; Thompson, C. B. Intracellular α-Ketoglutarate Maintains the Pluripotency of Embryonic Stem Cells. *Nature* **2015**, *518* (7539), 413–416.
(56) Wong, C. C.; Qian, Y.; Yu, J. Interplay between Epigenetics and Metabolism in Oncogenesis: Mechanisms and Therapeutic Approaches. *Oncogene* **2017**, *36* (24), 3359–3374.

(57) Wu, D.; Hu, D.; Chen, H.; Shi, G.; Fetahu, I. S.; Wu, F.; Rabidou, K.; Fang, R.; Tan, L.; Xu, S.; Liu, H.; Argueta, C.; Zhang, L.; Mao, F.; Yan, G.; Chen, J.; Dong, Z.; Lv, R.; Xu, Y.; Wang, M.; Ye, Y.; Zhang, S.; Duquette, D.; Geng, S.; Yin, C.; Lian, C. G.; Murphy, G. F.; Adler, G. K.; Garg, R.; Lynch, L.; Yang, P.; Li, Y.; Lan, F.; Fan, J.; Shi, Y.; Shi, Y. G. Glucose-Regulated Phosphorylation of TET2 by AMPK Reveals a Pathway Linking Diabetes to Cancer. *Nature* **2018**, *559* (7715), 637–641.
(58) Wellen, K. E.; Hatzivassiliou, G.; Sachdeva, U. M.; Bui, T. V.; Cross, J. R.; Thompson, C. B. ATP-Citrate Lyase Links Cellular Metabolism to Histone Acetylation. *Science* **2009**, *324* (5930), 1076–1080.
(59) Institute of Medicine; Food and Nutrition Board; Subcommittee on Upper Reference Levels of Nutrients; Standing Committee on the Scientific Evaluation of Dietary Reference Intakes and its Panel on Folate, Other B Vitamins, and Choline. *Dietary Reference Intakes for Thiamin, Riboflavin, Niacin, Vitamin B6, Folate, Vitamin B12, Pantothenic Acid, Biotin, and Choline*; National Academies Press, 2000.
(60) Myers, S. A.; Panning, B.; Burlingame, A. L. Polycomb Repressive Complex 2 Is Necessary for the Normal Site-Specific O-GlcNAc Distribution in Mouse Embryonic Stem Cells. *Proc. Natl. Acad. Sci. U. S. A.* **2011**, *108* (23), 9490–9495.
(61) Olivier-Van Stichelen, S.; Hanover, J. A. You Are What You Eat: O-Linked N-Acetylglucosamine in Disease, Development and Epigenetics. *Curr. Opin. Clin. Nutr. Metab. Care* **2015**, *18* (4), 339–345.
(62) Xu, Y.; Wang, Y.; Yan, S.; Zhou, Y.; Yang, Q.; Pan, Y.; Zeng, X.; An, X.; Liu, Z.; Wang, L.; Xu, J.; Cao, Y.; Fulton, D. J.; Weintraub, N. L.; Bagi, Z.; Hoda, M. N.; Wang, X.; Li, Q.; Hong, M.; Jiang, X.; Boison, D.; Weber, C.; Wu, C.; Huo, Y. Intracellular Adenosine Regulates Epigenetic Programming in Endothelial Cells to Promote Angiogenesis. *EMBO Mol. Med.* **2017**, *9* (9), 1263–1278.
(63) Gongol, B.; Sari, I.; Bryant, T.; Rosete, G.; Marin, T. AMPK: An Epigenetic Landscape Modulator. *Int. J. Mol. Sci.* **2018**, *19* (10). https://doi.org/10.3390/ijms19103238.
(64) Sakabe, K.; Wang, Z.; Hart, G. W. β-N-Acetylglucosamine (O-GlcNAc) Is Part of the Histone Code. *Proceedings of the National Academy of Sciences* **2010**, *107* (46), 19915–19920.
(65) Hottiger, M. O. Nuclear ADP-Ribosylation and Its Role in Chromatin Plasticity, Cell Differentiation, and Epigenetics. *Annu. Rev. Biochem.* **2015**, *84*, 227–263.
(66) Kok, D. E.; Steegenga, W. T.; McKay, J. A. Folate and Epigenetics: Why We Should Not Forget Bacterial Biosynthesis. *Epigenomics* **2018**, *10* (9), 1147–1150.
(67) Boon, R.; Silveira, G. G.; Mostoslavsky, R. Nuclear Metabolism and the Regulation of the Epigenome. *Nat Metab* **2020**, *2* (11), 1190–1203.
(68) Liu, W. H.; Zheng, J.; Feldman, J. L.; Klein, M. A.; Kuznetsov, V. I.; Peterson, C. L.; Griffin, P. R.; Denu, J. M. Multivalent Interactions Drive Nucleosome Binding and Efficient Chromatin Deacetylation by SIRT6. *Nat. Commun.* **2020**, *11* (1), 5244.
(69) van der Knaap, J. A.; Verrijzer, C. P. Undercover: Gene Control by Metabolites and Metabolic Enzymes. *Genes Dev.* **2016**, *30* (21), 2345–2369.
(70) Maher, M.; Diesch, J.; Casquero, R.; Buschbeck, M. Epigenetic-Transcriptional Regulation of Fatty Acid Metabolism and Its Alterations in Leukaemia. *Front. Genet.* **2018**, *9*, 405.
